# Structural insights into glycine reuptake inhibition

**DOI:** 10.1101/2020.12.20.110478

**Authors:** Azadeh Shahsavar, Peter Stohler, Gleb Bourenkov, Iwan Zimmermann, Martin Siegrist, Wolfgang Guba, Emmanuel Pinard, Markus A. Seeger, Thomas R. Schneider, Roger J.P. Dawson, Poul Nissen

## Abstract

The human glycine transporter 1 (GlyT1) regulates glycine mediated neuronal excitation and inhibition through sodium- and chloride-dependent reuptake of the neurotransmitter^1-3^. Inhibition of glycine reuptake via GlyT1 prolongs neurotransmitter signaling and has long served as a key therapeutic development strategy for treatment of a broad range of central nervous system disorders including schizophrenia and cognitive impairments^4^. Using an inhibition state-selective sybody and serial synchrotron crystallography, we determined the structure of GlyT1 in complex with a benzoylpiperazine chemotype inhibitor at 3.4 Å resolution. The inhibitor locks GlyT1 in an inward-open conformation and binds at the intracellular gate of the release pathway, overlapping with the glycine release site. The inhibitor likely reaches GlyT1 from the cytoplasmic leaflet of the plasma membrane. The study defines the mechanism of non-competitive inhibition and enables the rational design of new, clinically efficacious GlyT1 inhibitors.

## Main

Glycine is a conditionally essential amino acid with a dual role in the central nervous system (CNS). It acts as a classical neurotransmitter at inhibitory glycinergic synapses, where it induces hyperpolarizing chloride influx at postsynaptic terminals through ionotropic glycine receptors^1,2^. Yet, as the obligatory co-agonist of the N-methyl-d-aspartate (NMDA) receptor, glycine positively modulates calcium-dependent neuronal excitation and plasticity at glutamatergic synapses^1,3^. Glycine homeostasis is tightly regulated by reuptake transporters, including glycine-specific GlyT1 and GlyT2, that belong to the secondary active neurotransmitter/sodium symporters (NSSs) of the solute carrier 6 (SLC6) transport family^5^. GlyT1 (*SLC6A9*) and GlyT2 (*SLC6A5*) share a sequence identity of approximately 50%, similar to other members of the NSS family, such as serotonin transporter (SERT), dopamine transporter (DAT) and gamma-aminobutyric acid (GABA) transporter (GAT). GlyT1 is located on presynaptic neurons and astrocytes surrounding both inhibitory glycinergic and excitatory glutamatergic synapses and is considered the main regulator of extracellular levels of glycine in the brain^1,6^.

At glutamatergic synapses, GlyT1 plays a key role by maintaining sub-saturating concentrations of regulatory glycine for the NMDA receptor^7,8^. Hypofunction of NMDA receptor is implicated in pathophysiology of schizophrenia^9^, but pharmacological interventions to directly enhance NMDA receptor neurotransmission in schizophrenic patients have been unsuccessful^10,11^. Selective inhibition of glycine reuptake by GlyT1 is an alternative indirect approach to elevate endogenous, extracellular levels of glycine and potentiate NMDA transmission^1,4^. Several chemotypes of potent and selective GlyT1 inhibitors such as Bitopertin have been developed to achieve antipsychotic and pro-cognitive activity treatment of schizophrenia^4,12^. Bitopertin showed clear signs of neuroplasticity enhancement^13,14^ via the NMDA receptor glycine site, however, failed to show efficacy in phase III clinical trials (at a reduced dose), and a drug candidate targeting GlyT1 has yet to emerge.

Studies of NSS and homologues have revealed an alternating-access mechanism^15^, which involves a Na^+^- (and Cl^−^- in eukaryotic NSS) gradient dependent binding and occlusion of the extracellular substrate and ions followed by a rearrangement to inward-facing state and subsequent intracellular opening and release of bound ions and substrate. The conformational rearrangements of transmembrane helices during the transport cycle expose the substrate binding site to either side of the membrane^16-23^. Bitopertin behaves functionally as a non-competitive inhibitor of glycine reuptake^24^, nevertheless, detailed structural information of the inhibitor binding site, selectivity and the underlying molecular mechanism of glycine reuptake inhibition have remained elusive. Here, we present the first structure of a glycine transporter, the human GlyT1 in complex with a highly selective Bitopertin-analogue^25,26^, Cmpd1, and an inhibition-state-selective synthetic nanobody (sybody). Cmpd1 has been patented as a more potent inhibitor targeting GlyT1 that contains a benzoylisoindoline scaffold originating from the Bitopertin chemical series^26^. The GlyT1 structure reveals the molecular determinants and mechanism of action for glycine reuptake inhibition.

### Stabilization and crystal structure determination of GlyT1

Wild-type human GlyT1 (encoded by the *SLC6A9* gene) is unstable when extracted from the membrane and contains unstructured termini and a large, flexible extracellular loop 2 (EL2). To enable structure determination, we screened for point mutations to increase thermal stability while preserving ligand-binding activity. For the final crystallization construct, we combined Leu153Ala, Ser297Ala, Ile368Ala and Cys633Ala point mutations, truncated N- and C-termini (Δ1-90/Δ685-706) and shortened EL2 (Δ240-256) (Methods). The addition of selective GlyT1 inhibitor Cmpd1 increases the thermal stability of the transporter further by 30.5°C (Fig. 1a). Indicative of high-affinity binding with a stabilizing effect, we measured a half maximal inhibitory concentration (IC_50_) for Cmpd1 of 12.5 and 7 nM on human and mouse GlyT1, respectively, (Fig. 1b) in a membrane-based competition assay with [^3^H]Org24598 compound^27^. We therefore purified GlyT1 in the presence of Cmpd1 and generated sybodies to further stabilize the transporter in this inhibition-state conformation and identified sybody Sb_GlyT1#7 binding to GlyT1 with an affinity of 9 nM^28^. Microcrystals of GlyT1 in complex with Sb_GlyT1#7 and inhibitor Cpmd1 were obtained in lipidic cubic phase. Merging oscillation patterns collected from 409 mounted loops containing micro crystals by a serial synchrotron crystallography approach yielded a complete data set at 3.4 Å resolution. The structure was determined by molecular replacement using the structures of inward-occluded bacterial multi-hydrophobic amino acid transporter (MhsT, PDB ID 4US3) and inward-open human SERT (PDB ID 6DZZ)^17,19^. A high quality of resulting electron density maps enabled us to unambiguously model human GlyT1 in complex with the sybody and bound ligand (Fig. 1c and Extended Data Fig. 1).

**Fig. 1.**
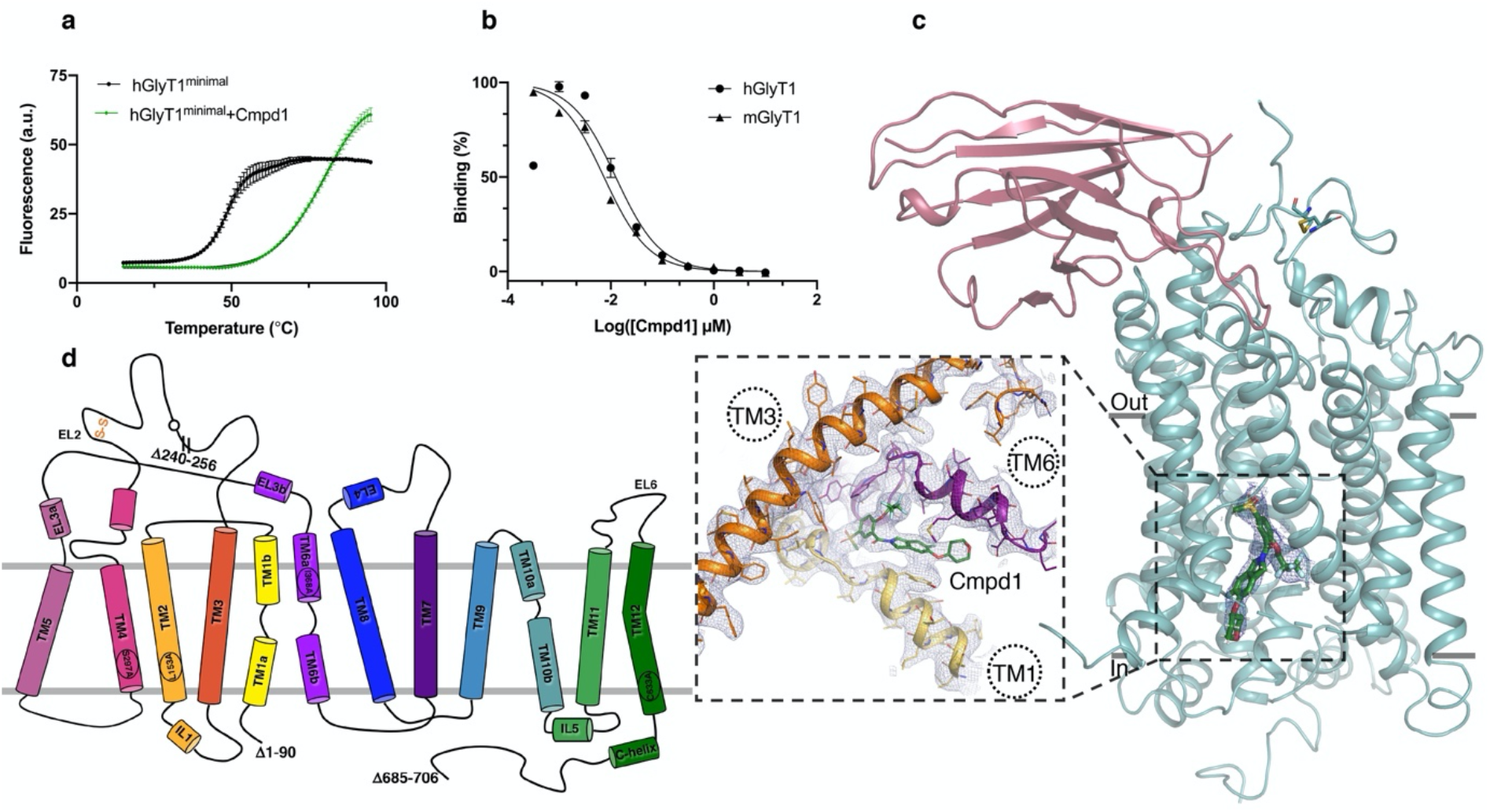
Stabilization, binding and recognition of inhibitor Cmpd1 by human GlyT1. (**a**) Thermal shift assays verify binding of Cmpd1 to GlyT1. Increasing concentrations of Cmpd1 show a strong dose-dependent stabilization on GlyT1 by raising the melting point from 48.8 to 79.3°C (mean ± SEM from quadruplicate measurements). GlyT1^minimal^ (containing N- and C-termini deletions) data with and without addition of the inhibitor is depicted in green and black, respectively. (**b**) Cmpd1 shows an IC_50_ of 7 and 12.5 nM on mouse and human GlyT1, respectively in membrane-based competition assays with radioactively labeled organon ([^3^H]Org24598). (**c**) Overall structure of human GlyT1 bound to selective inhibitor Cmpd1 and inhibition-state selective sybody. The magnified view of inhibitor binding pocket in 2*F*o – *F*c electron density map (blue) countered at 1.0 r.m.s.d. is depicted. TM8 is not shown for clarity. (**d**) Topology diagram of GlyT1 crystallization construct. The first 90 and last 31 residues have been removed from the N- and C-termini, respectively. EL2 is truncated from residue 240 to 256. The one remaining glycosylation site at Asn237 is shown as a sphere on EL2. The conserved disulfide bridge between Cys220 and Cys229 is shown on EL2. Single point mutations to improve thermal stability of the crystallization construct are shown on the respective transmembrane helices.

### GlyT1 architecture and conformation

GlyT1 (*SLC6A9*) adopts the general SLC6 transporters architecture of 12 alpha-helical transmembrane segments (TMs 1-12) with an inverted pseudo-two-fold symmetric architecture relating two transmembrane domains, TMs 1-5 and 6-10, denoted as the LeuT fold^17,18,21,22,29^(Fig. 1c, d). The transporter structure exhibits an inward-open conformation and superposition of GlyT1 to that of inward-open structures of SERT and leucine transporter (LeuT) and inward-oriented occluded MhsT yields C*α* root mean square deviations of 1.8, 2.3 and 3.2 Å, respectively (for details see Methods). TM1 and TM6 possess non-helical segments in the middle of the lipid bilayer, which coordinate Na^+^ and Cl^−^ ions^18,20,21^, accommodate substrates and inhibitors of various sizes^18,19,22^, and stabilize the ligand-free return state^17^. The intracellular part of TM5 is unwound at the conserved helix-breaking GlyX_9_Pro motif^17^ (Gly_313_X_9_Pro_323_ in GlyT1), and the N-terminal segment of TM1 (TM1a) is bent away from the core of GlyT1, which open the intracellular pathway to the center of the transporter (Fig 2a). The splayed motion of TM1 disrupts the interaction between the conserved residues Trp103 of TM1a and Tyr385 at the cytoplasmic part of TM6 that is otherwise present in outward-open and occluded conformations (Extended Data Figs. 2, 3)^17,18,20,22^.

**Fig. 2.**
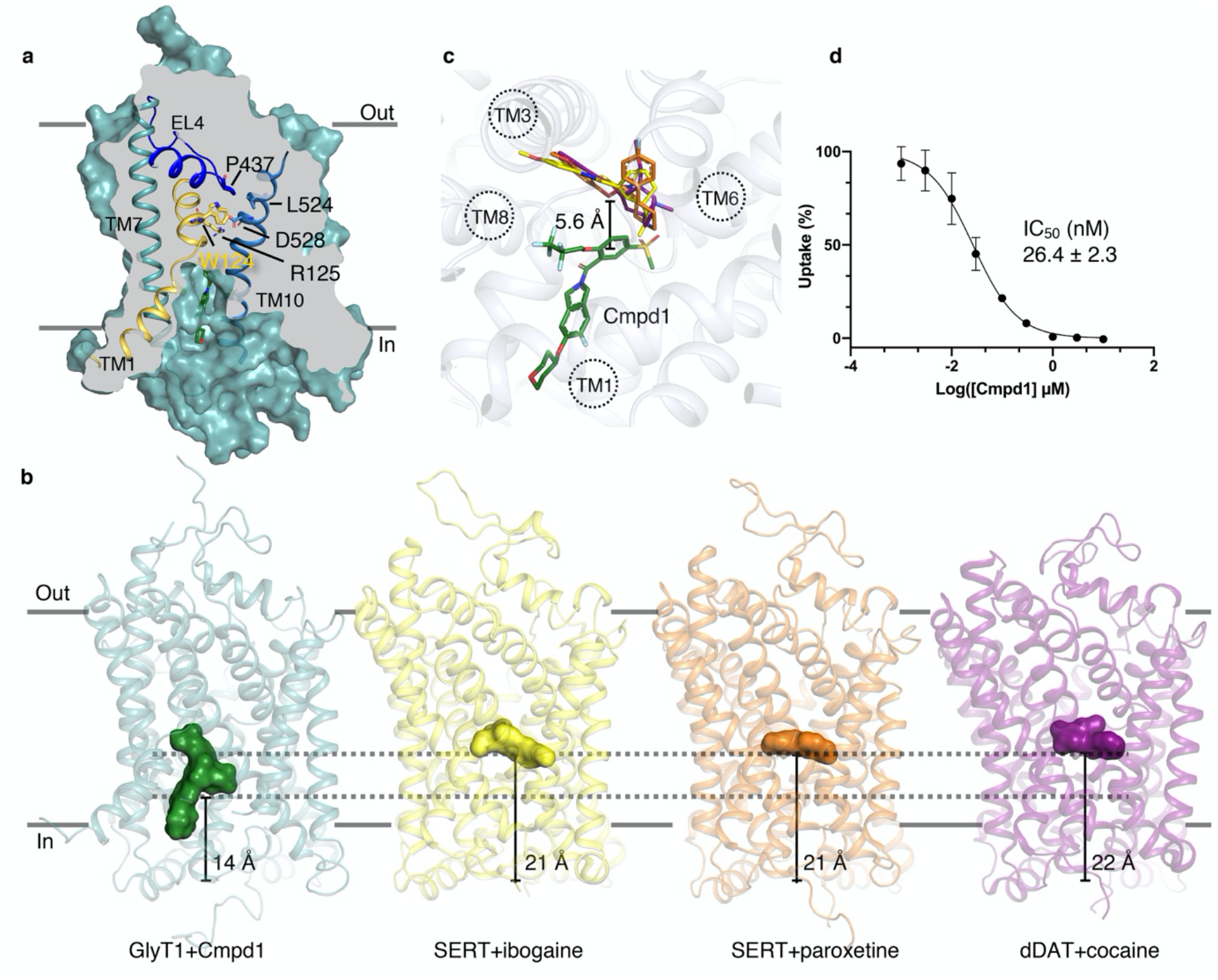
Uptake inhibition and binding mode of Cmpd1 at inward-open GlyT1. (**a**) Surface representation of the GlyT1 inward-open structure viewed parallel to the membrane. Closed extracellular vestibule around Trp124 (yellow) and open intracellular pathway are displayed. Residues R125 (TM1), P437 (EL4), L524, and D528 (TM10) are shown as sticks. (**b**) and (**c**) Binding mode comparison of Cmpd1 in GlyT1 with inhibitor binding site at other NSS transporters. Cmpd1 bound at the proximity of the central binding site in GlyT1 is colored in green. Paroxetine (orange) and ibogaine (yellow) bound at the central binding site of SERT (PDB IDs 5I6X and 6DZY, respectively) and cocaine (purple) bound to dDAT (PDB ID 4XP4) are shown as examples. The difference in location of the bound ligands in transporters is marked with dotted lines in (**b**). (**c**) Compared to paroxetine, ibogaine and cocaine, Cmpd1 is located 5.6 ± 0.1 Å further away from the center of the transporter. The distance is measured between the center of the phenyl ring of Cmpd1 and center of mass of other NSS inhibitors shown in the figure. (**d**) Cmpd1 inhibits the uptake of glycine by human GlyT1 with an IC_50_ of 26.4 nM.

Comparison of GlyT1 with inward-open SERT (PDB ID 6DZZ) shows structural differences mainly at the intracellular halves of the helices (Extended Data Fig. 4a-e), and in particular at the intracellular gate of GlyT1 defined by TM1a and TM5. The intracellular half of TM5 has splayed away by 17°, whereas TM1a is by 29° closer to the transporter core compared to corresponding segments of SERT (Extended Data Fig. 4b). As a result, the intracellular gate, measured as C*α*-C*α* distance between conserved Trp103 on TM1a and Val315 on TM5, is by 4 Å more closed than that of the inward-open structure of ibogaine-bound SERT (Extended Data Fig. 4e). On the extracellular side, a C*α*-C*α* distance of 8.9 Å between Arg125 of TM1a and Asp528 of TM10, and a close packing of the extracellular vestibule around Trp124 in the conserved NVWRFPY motif of TM1 indicates a closed extracellular gate (Fig. 2a, Extended Data Figs. 2, 3).

Intracellular loops (ILs) 1 to 5 together with the unwound region of TM5 and a C-terminal helix form the cytoplasmic interface. The C-terminal tail of the transporter caps over the intracellular face stabilized by interactions with IL1 and IL5 (Extended Data Fig. 3). Similar to dDAT and hSERT, TM12 of GlyT1 kinks at Ser_620_ of the Gly_613_(X_6_)Ser(X_4_)Pro_625_ motif conserved in eukaryotic NSS transporters (Extended Data Figs. 1-3).

The extracellular part of the structure is mainly composed of a long extracellular loop 2 (shortened in this structure), EL3, EL4 and EL6. EL2 carries a strictly conserved disulfide bridge (Cys220-Cys229) and four N-linked glycosylation sites, Asn237, Asn240, Asn250 and Asn256 (Fig. 1d, Extended Data Figs. 1-3). Three glycosylation sites were removed by the EL2 truncation, but Asn237 was critical for ligand binding likely through correct trafficking of the transporter to the plasma membrane^30^.

The conformation-specific sybody binds through several interactions to the extracellular segment of GlyT1 involving EL2, EL4, TM5 and TM7 (Fig. 1c, Extended Data Fig. 5). Sb_GlyT1#7 is selective for the inward-open conformation of GlyT1 and shows a conformation-stabilizing effect as evidenced by an increase of 10°C in thermal stability and an apparent affinity increase for [^3^H]Org24598 of almost 2-fold in a scintillation proximity assay^28^. In the crystal, sybody has a central role in formation of lattice contacts, packing against the neighboring sybody (Extended Data Fig. 6).

### Unique NSS transporter binding mode

Unambiguous density for the inhibitor Cmpd1 was observed in proximity of the GlyT1 central binding pocket between the transmembrane helices 1, 3, 6 and 8 (Fig. 1c and Extended Data Fig. 7). Comparison of the inhibitor binding site in GlyT1 with the equivalent site of other NSS structures shows that Cmpd1 is in 6.0 ± 0.5 Å distance from the core with its center of mass located 14 Å above the cytosolic end of the transporter while inhibitors of SERT and dDAT bind at the central binding site within 21-22 Å of the cytosolic face (Fig. 2b, c). Furthermore, the inhibitor binds GlyT1 in a unique binding mode, lodged in proximity of the transporter’s center and extending into the intracellular release pathway of substrate and ions between TM6b and TM1a accessible to solvent. This mode of inhibition is not observed in other NSS-inhibitor complexes (Fig. 2b, c).

Cmpd1 is from the benzoylisoindoline class of selective GlyT1 inhibitors^25^ that inhibits the uptake of glycine in mammalian (Flp-in™-CHO) cells, expressing mouse^26^ or human GlyT1 with an IC_50_ of 26.4 and 7 nM, respectively (Fig. 2d). The isoindoline scaffold of Cmpd1 forms a *π*-stacking interaction with Tyr116 of TM1 (Fig 3a, b). The phenyl ring is engaged in an edge-to-face stacking interaction with the aromatic ring of Trp376 located on the unwound region of TM6. The inhibitor is further stabilized by hydrogen bond and van der Waals interactions with residues from TM1, TM3, TM6 and TM8. (Fig. 3a, b).

**Fig. 3.**
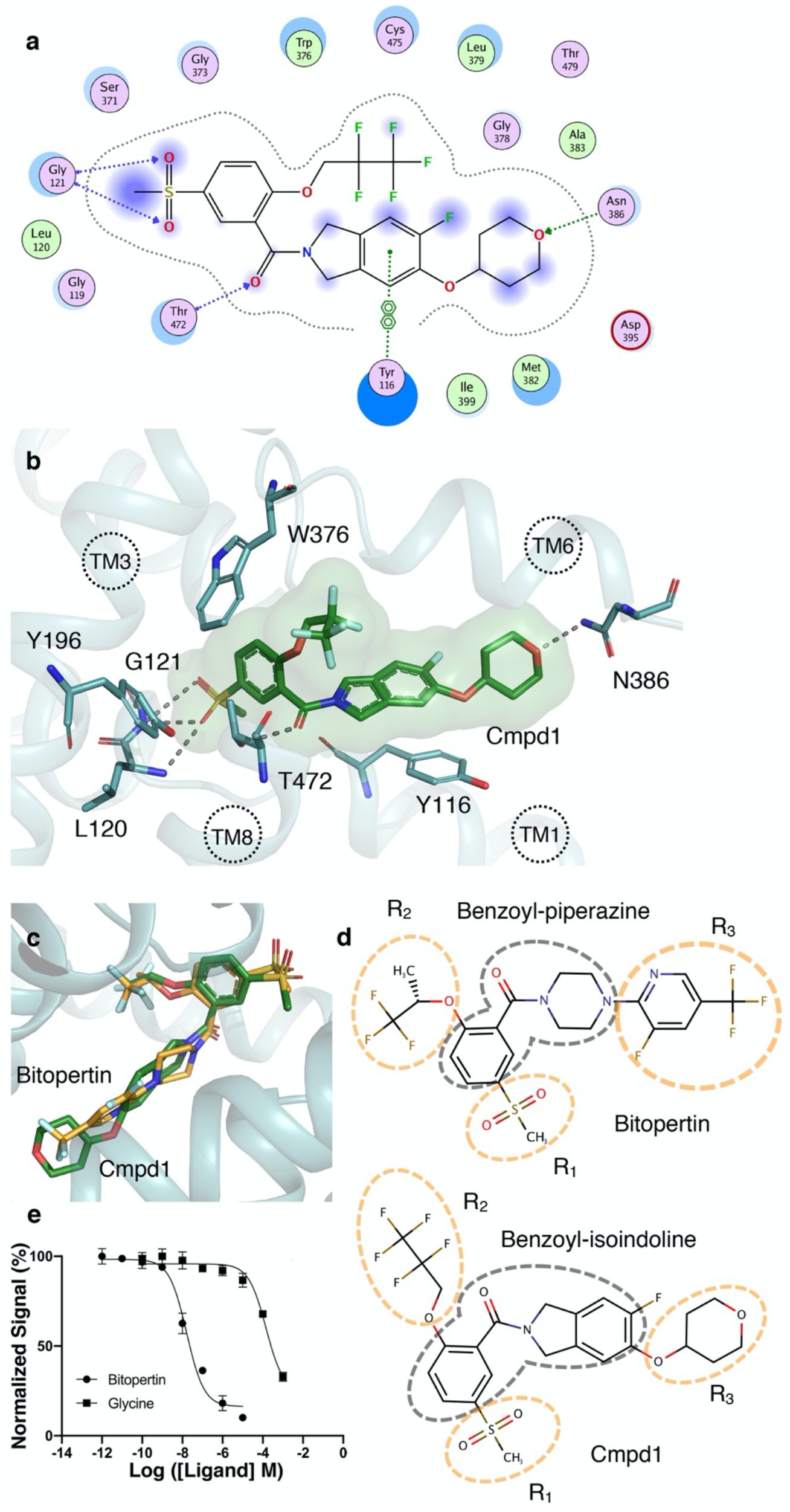
Binding pocket. (**a**) Protein-ligand interactions diagram calculated with MOE. Several hydrogen bonds that contribute to ligand binding are shown with dotted lines (backbone interaction in blue and side chain interactions in green). The π-stacking interaction of the isoindoline scaffold of the ligand and Tyr116 is depicted. Hydrophilic residues are shown in purple, blue rings indicate basic groups, red rings indicate acidic groups and hydrophobic residues are depicted in green. (**b**) Close-up view of Cmpd1 binding pocket at GlyT1. The two ends of the inhibitor are stabilized by hydrogen bond interactions with residues from TM1 and TM6; the backbone amine of Gly121 and Leu120 hydrogen bond to sulfonyl oxygen atoms and Asn386 from TM6 to the oxygen atom of the tetrahydropyran moiety of the inhibitor. From TM8, the hydroxyl group of Thr472 participates in a hydrogen bonding interaction with the carbonyl oxygen of the scaffold as well. The aromatic ring of Tyr116 localized 4.2 Å from isoindoline scaffold of the compound (π-stacking interaction) is shown. Hydroxyl group of Tyr196 from TM3 is likely in a weaker hydrogen bond interaction with methyl-sulfone moiety of the inhibitor. Inhibitor binding is also supported by an edge-to-face stacking interaction between the phenyl ring of the ligand and the aromatic sidechain of Trp376. (**c**) Docking of Bitopertin (orange) into the GlyT1 inhibitor-binding pocket showing Bitopertin’s binding mode compared to Cmpd1 (green). (**d**) Comparison of Bitopertin (benzoylpiperazine series, top) and Cmpd1 (benzoylisoindoline series, bottom). The scaffolds of the compounds are marked with a gray dashed line and the three R groups are marked with orange dashed lines. (**e**) Scintillation proximity competition assays using [^3^H]Org24598 and varying concentrations of Bitopertin and glycine showing that Bitopertin and glycine also compete for binding at GlyT1.

We generated a stable construct with the single point mutation Ile192Ala that was not able to bind the inhibitor (Extended Data Fig. 8). Interestingly, Ile192 is within van der Waals distance of the Trp376 side chain, which is stabilized in a rotamer perpendicular to the phenyl ring of the inhibitor (Extended Data Fig. 8). Trp376 is the bulky hydrophobic residue of a conserved (G/A/C)ΦG motif in the unwound segment of TM6 that determines the substrate selectivity of SLC6 transporters^31,32^, and the AWG sequence observed in GlyT1 is indeed fitting for the small glycine substrate. Ile192, although not in direct interaction with the inhibitor, plays an important role for binding of Cmpd1 through reduced freedom of the Trp376 side chain, which also further restricts the binding pocket for glycine.

Adding a Lichenase fusion protein construct (PDB ID 2CIT^33^) to the N-terminus of the GlyT1 construct, we generated and crystallized also a GlyT1-Lic fusion protein in complex with Sb_GlyT1#7 and obtained a 3.9 Å resolution data set collected from 1222 mounted loops containing micro crystals (Extended Data Fig. 6b). The electrogenic reuptake of glycine via GlyT1 is coupled to the transport of two Na^+^ and one Cl^−^ ions. Both GlyT1 and GlyT1-Lic constructs were purified and crystallized in the presence of 150 mM NaCl, yet, we observe no electron density for Na^+^ or Cl^−^ ions in the GlyT1 structure. However, the Cl^−^ and Na^+^ binding sites were evident in the lower resolution map of the GlyT1-Lic crystal structure, which may have captured a preceding state in transitions associated with ion release to the intracellular environment (Extended Data Fig. 9).

### Plasticity of the binding pocket

Similar to reported benzoylisoindolines^25^, Cmpd1 is >1000-fold selective for GlyT1 against GlyT2 (Fig. 1b and Extended Data Fig. 10). Comparing the binding pocket residues of GlyT1 to corresponding residues in GlyT2 points to direct clues. Gly373 in GyT1 corresponds to Ser497 in GlyT2. Interestingly, N-methyl glycine (sarcosine) and N-ethyl glycine are substrates of GlyT1 and the Ser497Gly mutant of GlyT2, but not of wild-type GlyT2^32,34,35^, which can be explained readily by a steric clash with Ser497. Furthermore, GlyT1 residues Met382 and Ile399 correspond to Leu and Val in GlyT2, respectively that diminish the van der Waals interactions between the inhibitor and the transporter.

Molecular docking places Bitopertin in the binding pocket of GlyT1 with its benzoylpiperazine scaffold matching the benzoylisoindoline scaffold of Cmpd1 (Fig. 3c). The binding mode and scaffold substituent interactions (R1 through R3) are supported by the previously reported structure-activity relationships of the benzoylpiperazine and benzoylisoindoline series^12,25^. The R1 pocket (methyl-sulfone moiety) is spatially constrained and prefers small, polar substituents with an H-bond acceptor group. The pocket harboring R2 substituents (O-C_3_F_5_) is mainly hydrophobic and accommodates linear and cyclic substituents up to a ring size of 6. The R3 (tetrahydropyran) pocket is fairly large, exposed to solvent and can accommodate diverse groups with different functionalities (Fig. 3d). We observed a higher flexibility for the tetrahydropyran moiety as the corresponding portion of the electron density was not well resolved. Considering the size and solvent exposure of this pocket, the R3 position is the favorable handle to fine-tune the properties of the inhibitor.

Superposition of glycine-bound LeuT (PDB ID 3F4J) and tryptophan-bound MhsT (PDB ID 4US3) on inhibitor-bound GlyT1 shows that the sulfonyl moiety of the inhibitor likely mimics the carboxyl group of the glycine substrate interacting with TM1 and TM3 (Extended Data Fig. 11)^17,36^. We observe that in 0.1 mM glycine concentration or higher, selective inhibitors of GlyT1 are competed out, further supporting overlapping binding sites (Fig. 3e).

### Mechanism of inhibition

Cmpd1, Bitopertin and similar inhibitors of GlyT1 were shown to be non-competitive glycine reuptake inhibitors^4,24^, yet, the binding site appears to overlap with the glycine substrate. Likely Cmpd1, Bitopertin and related chemotypes diffuse across the cell membrane and bind from the cytoplasmic side to an inward-open structure that involves unwinding the TM5 segment and a hinge-like motion of TM1a to fit the bulky inhibitor (Fig. 4). Glycine on the other hand, binds with a high-affinity to the outward-open conformation, which is at the same time exposed to high concentrations of the driving Na^+^ and Cl^−^ ions at the synaptic environment. Following binding of glycine and ions, the transporter transforms to an inward-open conformation, exerting low glycine affinity. Release of ions and glycine from the inward-open state is essentially irreversible, enabling non-competitive inhibitors of transport to bind and shift the conformational equilibrium towards an inward-open conformation. (Fig. 4). Similar to ibogaine inhibition of SERT^37^, the binding sites of glycine and non-competitive inhibitors of GlyT1 explore two distinct conformational states, outward and inward oriented.

**Fig. 4.**
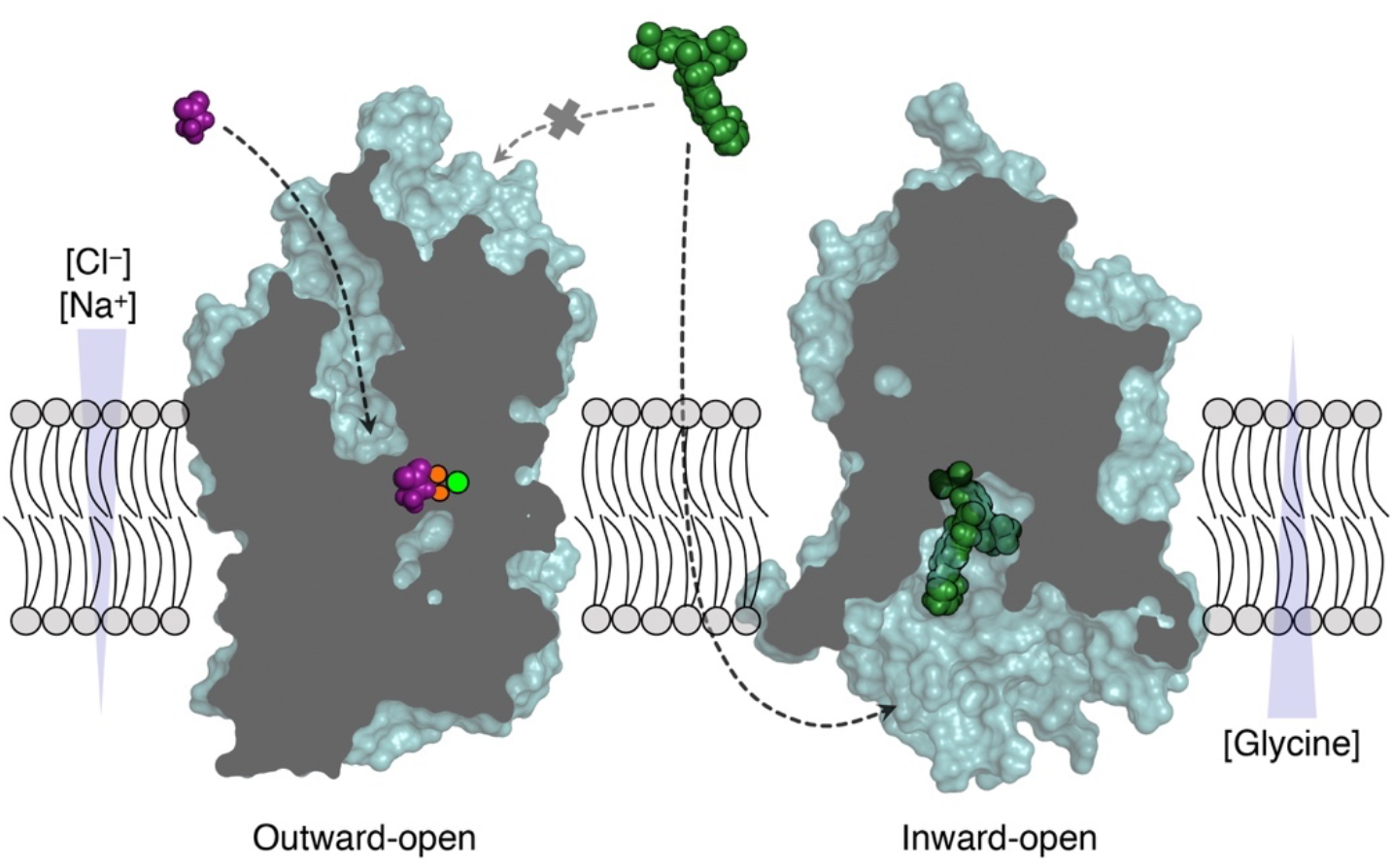
Mechanism of inhibition at GlyT1. Glycine (purple) binds with a high affinity to the outward-open conformation of GlyT1 (left, based on a dDAT homology model, PDB ID 4M48) that is exposed to high concentrations of the driving sodium and chloride ions (orange and green spheres, respectively) at the synaptic environment. The non-competitive inhibitor Cmpd1 (green) diffuses through the synaptic cell membrane and reaches the intracellular side of GlyT1. Cmpd1 locks the transporter in an inward-open conformation (right) with the characteristic hinge-like motion of TM1a and unwinding of TM5. Cmpd1 inhibits GlyT1 by shifting the conformational equilibrium to the inward-open state.

Considering the high membrane permeability measured for Bitopertin^12^, it is likely that the inhibitor dissipates into locations other than the synapse. In fact, GlyT1 is also expressed in peripheral tissues including erythrocytes where glycine plays a key role in heme biosynthesis. Inhibition of GlyT1 by Bitopertin in these cells results in a tolerable decreased in the level of hemoglobin. However, possible risks associated with such effect was a prohibitory factor in phase III clinical trial of Bitopertin, which was administered at a lower dose than the proof-of-concept phase II clinical studies. It further remains unclear if Bitopertin administration reached optimal GlyT1 occupancies in patients or a higher placebo response in clinical trials resulted in indistinguishable efficacy of Bitopertin over placebo^10,38,39^.

## Conclusion

Clinical trials of Bitopertin and other selective inhibitors of GlyT1 are challenging due to the complex nature of schizophrenia neurobiology and glycinergic signaling. Efforts for developing selective inhibitors can be re-evaluated in light of the GlyT1 structure and binding site described here.

The inhibitor-bound GlyT1 complex captures the transporter in an inward-open state and reveals a unique mode of NSS inhibitors. Binding pocket interactions explain the selectivity of the inhibitor against GlyT2. The structure reveals how a bulky inhibitor lodges between transmembrane helices in the middle of the transporter and extends into the intracellular release pathway for ions and substrate, where it is exposed to the cytosolic environment. The non-competitive mechanism of action occurs through shifting the conformational equilibrium from outward-open to inward-open by stabilizing the inward-open conformation of GlyT1 associated with release of ions and substrate.

The sybody Sb_GlyT1#7 too is highly selective for the inhibited, inward-open conformation of GlyT1. Recent efforts to engineer antibodies and achieve effective targeting and efficient crossing of the blood brain barrier^40^ to deliver an inhibition-state-specific sybody represents an alternative approach to small molecule inhibitors of GlyT1. The structure of human GlyT1 provides a platform for the rational design of new small molecule inhibitors and antibodies to develop selective inhibitors targeting the glycine reuptake transporter.

## Acknowledgments

We are grateful to Ralf Thoma (F. Hoffmann-La Roche) for long-term project support and fruitful discussions, and Lisa Hetemann and Jennifer Gera for their early contributions in producing the mutants. We thank Fabio Dall’Antonia for writing the ctrl-d script. We thank Joanna Pieprzyk for assistance in mammalian cell expression and Hanne Poulsen for assistance in electrophysiology studies. We thank Maria M. Garcia Alai for access to sample preparation and crystallization facility at EMBL, Hamburg and Ioana Maria Nemtanu and Christian Guenther for technical assistance. We acknowledge excellent support at P14 beamline operated by EMBL Hamburg at the PETRA III storage ring (DESY, Hamburg). We are grateful to Janet Thornton for advice and valuable discussions on the project, and to Joseph A. Lyons, Bjørn Pedersen and Christian Loew for valuable discussion. This project has received funding from Novo Nordisk Foundation (to P.N.), the Lundbeck Foundation via the DANDRITE neuroscience center, and F. Hoffmann-La Roche LTD. Azadeh Shahsavar was supported by EI3POD program fellowship under Marie Sklodowska Curie COFUND.

## Author contributions

R.J.P.D. initiated the project and designed the project with P.N. and T.R.S. at Roche, Aarhus University, and EMBL. Construct design was performed by R.J.P.D. and construct screening was done by P.S., A.S. and R.J.P.D. Expression was performed by P.S., M.S. and A.S. Purification was performed by P.S. and A.S. Radioligand and thermal-shift assays were performed by E.P. and P.S. Sybody was generated by I.Z., M.A.S., P.S. and R.J.P.D. Crystallization, data collection, processing and structure refinement were performed by A.S. Serial data collection was established by A.S., G.B. and T.R.S. Dozor and serial data processing scripts were modified by G.B. Molecular modeling and docking were done by W.G. Manuscript was written by A.S., R.J.P.D., and P.N. with contributions from T.R.S., G.B. and W.G.

## Data availability

Coordinates and structure factors of 3.4 Å structure of GlyT1 have been deposited in the Protein Data Bank under accession code 6ZBV and the second structure of GlyT1 bound to Na and Cl ions will be fully deposited before publication.

## Code availability

For statistical analysis of serial diffraction data, the program Ctrl-D version 0.3 build 465 as developed by EMBL-Hamburg was used. The linux-executable can be obtained from upon request.

## Methods

### GlyT1 constructs

Human GlyT1 cDNA sequence was codon optimized and synthesized by Genewiz for mammalian cell expression, and GlyT1-Lic for insect cell expression. Both constructs contain N- and C-termini deletions of residues 1-90 and 685-706, respectively as well as a deletion in the extracellular loop 2 (EL2) between residues 240-256. To improve thermal stability of the constructs, point mutations L153A, S297A, I368A and C633A have been introduced. In addition, the N-terminal residue 91 has been omitted from GlyT1-Lic sequence and residues 9-281 of Lichenase (PDB ID 2CIT) have been fused at the N-terminal. The sequences of GlyT1 and GlyT1-Lic followed by a C-terminal enhanced green fluorescent protein (eGFP) and a decahistidine-tag have been cloned into a pCDNA3.1 vector for transient transfection in human embryonic kidney (HEK293) cells and pFastBac vector for baculovirus expression in *Spodoptera frugiperda* (*Sf9*) insect cells, respectively.

### Transporter expression and purification

GlyT1 was expressed in FreeStyleTM 293 Expression Medium in 1 L scale in 600 mL TubeSpin® Bioreactors incubating in the orbital shaker at 37°C, 8% CO2, 220 RPM in humidified atmosphere. The cells were transfected at the density of 1 × 10^6^ cells/mL and viability of > 95%. Linear polyethylenimine 25 kDa, (LPEI 25 K) was used as the transfection reagent with a GlyT1 DNA: LPEI ratio of 1:2. The cells were typically harvested 60 hours post transfection at a viability of around 70% and stored at −80°C until purification.

GlyT1-Lic was expressed in 20-25 L scale in 50 L single use WAVE bioreactors (CultiBag RM, sartorius stedim biotech) at 27°C with 18-25 rocks/min in a 40% oxygenated Sf900-III medium (Invitrogen). The cells were typically infected with a 0.25% volume of infection of the virus at a density of 2-3 × 10^6^ cells/mL and viability of > 95%. The cells were harvested 72 hours post infection at a viability of around 80% and stored at −80°C until purification.

Purification of GlyT1 constructs has been previously described in details^28^. In brief, the biomass was solubilized in 50 mM Tris-HCl pH 7.5, 150 mM NaCl, 100 µM Cmpd1 ([5-Methanesulfonyl-2-(2,2,3,3,3-pentafluoro-propoxy)-phenyl]-[5-tetrahydro-pyran-4-yloxy)- 1,3-dihydro-isoindol-2-yl]-methanone), 15-25 µM brain polar lipids extract (Avanti), containing either 1% (w/v) lauryl maltose neopentyl glycol (LMNG) or 1% (w/v) decyl maltose neopentyl glycol (DMNG) and 0.1% cholesteryl hemisuccinate (CHS). The protein was purified by batch purification using TALON affinity resin (GE Healthcare) and treated with HRV-3C protease (Novagen) to cleave the eGFP-His-tag and Roche PNGase F (*F. meningosepticum*) to trim glycosylation. The transporter was concentrated typically to 15-30 mg/mL in the final buffer containing 50 mM Tris-HCl pH 7.5, 150 mM NaCl, 50 µM inhibitor, 15-25 µM brain polar lipids extract, containing 0.01% LMNG (w/v) - 0.001% CHS for GlyT1 and either 0.05% (w/v) LMNG - 0.005% CHS or 0.01% DMNG - 0.001% CHS for GlyT1-Lic construct.

### Lipidic cubic phase crystallization

Prior to crystallization, the concentrated GlyT1 was incubated with Sb_GlyT1#7 in 1:1.2 molar ratio (GlyT1: sybody) and 1 mM inhibitor. The protein solution was reconstituted into mesophase using molten monoolein spiked with 5% (w/w) cholesterol (Sigma) in 2:3 ratio of protein solution: lipid using two coupled Hamilton syringes. Crystallization trials were carried out in 96-well glass sandwich plates (VWR) by a Gryphon LCP crystallization robot or a Mosquito LCP dispensing robot in a humidified chamber using 50-100 nL of mesophase overlaid with 800 nL of crystallization solution. The plates were incubated at 19.6°C and inspected manually. Crystals appeared in 3-10 days in 0.1M ADA pH 7, 13-25% PEG600, 4-14% v/v (±)-1,3-Butanediol with the longest dimension of 2-5 μm. The micrometer-sized crystals were harvested from the LCP matrix using MiTeGen MicroMounts and flash frozen in liquid nitrogen.

### Data collection, processing and structure determination

Crystallographic data were collected at P14 beamline operated by EMBL Hamburg at the PETRA III storage ring (DESY, Hamburg) using the 5 × 10 μm^2^ (Vertical × Horizontal) microfocus beam with a total photon flux of 1.3 × 10^13^ photons/second at the sample position. Diffraction data were recorded on an EIGER 16M detector. We employed a data collection strategy where we typically defined a region of interest of 60 × 14 – 290 × 340 µm^2^ on the loop containing crystals oriented perpendicular to the incoming beam. Diffraction data were collected using serial helical line scans with 1 μm sample displacement along the rotation axis during the acquisition of one frame and an oscillation of 0.2° at an exposure time of 0.1 s with 100% transmission.

Dozor was used as the first step of data processing to identify diffraction patterns within the large set of frames. Each diffraction image was analyzed by Dozor^41,42^, which determined a list of coordinates for diffraction spots and their partial intensities, followed by generation of a diffraction heat map.

Diffraction data was indexed and integrated using XDS^43,44^ and resulting partial mini data sets containing 3-20 consecutive images were scaled with XSCALE^44^. In some cases, mini-data sets adjacent in the frame number were merged into longer data sets (>20 frames) manually. One rotation data set of 20 frames with oscillation of 1.0° is included in GlyT1-Lic data set.

The choice of partial mini data sets to be merged into a high-quality complete data set was guided by an inhouse script, ctrl-d, that measured the correlation of each mini data set to the rest of mini data sets. The important criterion was the requirement of enough number of collected data sets to have a scaling model for robust estimation of outliers.

A total of 514 2D helical scans were performed on 409 mounted loops containing micro crystals of GlyT1 that resulted in collection of 1365232 diffraction patterns of which 30837 frames contained more than 15 diffraction spots. 229 mini data sets were indexed and integrated of which 207 mini data sets, containing 3400 frames, with correlation of above 0.7 were scaled and merged (Extended Data Table 1, Extended Data Fig. 12a). For GlyT1-Lic, a total number of 1733 2D helical scans were performed on 1222 mounted loops containing micro crystals that resulted in collection of 3190397 diffraction images of which 225037 contained equal or more than 15 spots. 249 mini data sets were indexed and integrated of which 213 mini data sets, containing 3906 diffraction patterns, with correlation of above 0.5 were scaled and merged (Extended Data Table 2, Extended Data Fig. 12b).

The structure of GlyT1-sybody complex was solved by molecular replacement using modified models of MhsT (PDB ID 4US3), and SERT (PDB ID 6DZZ), where the loops, TM12 and C-terminal tail were removed from the original models and an ASC binding nanobody (PDB ID 5H8D) as separate search models in Phaser. To solve the structure of GlyT1-Lic the Lichenase fusion protein structure (PDB ID 2CIT) was used as the third search model. The models were refined with Buster followed by visual examination and manual rebuilding in *Coot* and ISOLDE^45-47^. The final data and refinement statistics are presented in Extended Data Table 3.

### Scintillation proximity assay

Scintillation proximity assays (SPA) were performed in 96-well plates (Optiplate, Perkin Elmer) using Copper HIS-Tag YSi SPA beads (Perkin Elmer) and radioactively labeled Organon ([^3^H]Org24598, 80Ci/mmole). Reactions took place in the assay buffer containing 50 mM TRIS pH 7.5, 150 mM NaCl and 0.001% LMNG supplemented with solubilized GlyT1 cell membrane/SPA mix (0.3 mg/well) and for competition experiments serially diluted non-labeled inhibitor Cmpd1 (0.001 nM to 10 μM final concentration), Bitopertin (0.001 nM to 10 μM), or Glycine (0.1 nM to 1 mM). Assays were incubated for 1 hour at 4°C before values were read out using a top count scintillation counter at room temperature. In thermal shift scintillation proximity assays (SPA-TS), solubilized protein was incubated for 10 min with a temperature gradient of 23-53°C across the wells in a Techne Prime Elite thermocycler before mixing with SPA beads.

### Fluorescence-detection size-exclusion chromatography-based thermostability assay (FSEC-TS)

FSEC-TS method was used to evaluate thermostability of constructs^48^. 180 μL aliquots of solubilized GlyT1 cell membrane were dispensed in a 4°C-cooled 96-well PCR plate (Eppendorf) in triplicates. A gradient of 30-54°C for 10 min was applied on the plate in a Bio-Rad Dyad thermal cycler. The plate was cooled down on ice and 40 μl of the samples were injected on a 300 mm Sepax column in 50 mM TRIS pH 7.5, 150 mM NaCl and 0.001% LMNG and the SEC profile was monitored using the eGFP-tag fluorescence signal.

### Thermofluor stability assay

A GlyT1^minimal^ construct (containing N- and C-termini deletions of residues 1-90 and 685-706, respectively) was used for thermofluor stability assays which was based on wild type GlyT1 containing N- and C-termini deletions of residues 1-90 and 685-706, respectively and was expressed in *Sf9* insect cells and purified as described above. Purified GlyT1^minimal^ was diluted in 50 mM Tris pH 7.5, 150 mM NaCl and 0.001% LMNG to a final concentration of 0.73 μM and distributed into the wells of a 96 well PCR plate on ice. The inhibitor was added to the wells at a final concentration of 10 μM and a corresponding amount of DMSO was added to the control wells. The plate was sealed and incubated for 30 minutes on ice. A 1:40 (v/v) working solution of the CPM (N-[4-(7-diethylamino-4-methyl-3-coumarinyl)phenyl]maleimide) dye stock (4 mg/mL in DMSO) was prepared and 10 μL of this solution was added to 75 μL of protein sample in each well and mixed thoroughly. A published assay based on CPM dye was adapted to perform the stability tests^49^. The melting profiles were recorded using a real-time PCR machine (Rotor-Gene Q, QIAGEN) with temperature ramping from 15°C to 95°C at a heating rate of 0.2°C/s. The melting temperatures (Tm) were calculated from the point of inflection based on a fit to the Boltzmann equation.

### Molecular modeling

The 3D conformer generator Omega (OpenEye) was used to generate a conformational ensemble for Bitopertin. Each conformer was superimposed via ROCS (OpenEye)^50^ onto the receptor-bound conformation of Cmpd1 and the overlay was optimized with respect to 3D shape similarity. The highest scoring conformer was retained and energy-minimized within the binding pocket using MOE^51^. Docking was performed with the software GOLD^52^ from CCDC with default settings and the standard scoring function ChemPLP. An additional energy minimization within the binding pocket was performed with the 5 best docking poses.

Rapido was used for structure superpositions^53^. A total number of 513, 414, and 393 residues were used for alignment of SERT (PDB ID 6DZZ), LeuT (PDB ID 3TT3) and MhsT (PDB ID 4US3) structures on GlyT1. Residue ranges used for alignment were 104-224, 226-232, 259-306, 316-353, 357-388, 390-433, 438-489, 491-632, 636-652 of GlyT1 and 83-152, 154-204, 206-212, 222-239, 242-271, 281-318, 322-353, 355-398, 404-597, 600-616 of SERT in SERT-GlyT1 superposition, 115, 117-211, 215-219, 262-270, 272-278, 281, 288-307, 317-352, 354-374, 376-387, 390-421, 429-489, 496-519, 522-530, 532-559, 568-592 of GlyT1 and 21-68, 71-73, 76-80, 82-87, 90-123, 126-130, 141-156, 160, 166-185, 196-217, 222-240, 242-257, 259-270, 273-291, 293-305, 307-312, 318-372, 374-406, 408-435, 444-468 of LeuT in GlyT1-LeuT superposition and 119-173, 176-210, 264-271, 318-352, 358-422, 432-487, 532-554, 568-595, 493-517, 287-306 of GlyT1 and corresponding residues 28-82, 88-122, 134-141, 178-212, 218-282, 284-339, 389-411, 421-448, 343-367, 148-167 of MhsT in GlyT1-MhsT superposition.

### Glycine uptake inhibition assay

Glycine uptake inhibition assays were performed in quadruplicate and according to a method previously described^24^. In brief, mammalian Flp-in™-CHO cells were transfected with human and mouse GlyT1 and human GlyT2 cDNA and were plated at a density of 40,000 cells/well in complete F-12 medium 24 h before uptake assays. The medium was aspirated the next day and the cells were washed twice with uptake buffer containing 10 mM HEPES-Tris pH 7.4, 150 mM NaCl, 1 mM CaCl2, 2.5 mM KCl, 2.5 mM MgSO4 and 10 mM (+) D-glucose. The cells were incubated for 20 min at 22 °C with no inhibitor, 10 mM non-radioactive glycine, or a concentration range of the inhibitor to calculate IC_50_ value. A solution containing 25 μM non-radioactive glycine and 60 nM [^3^H]glycine (11-16 Ci/mmol) (hGlyT1 and mGlyT1) or 200 nM [^3^H]glycine (hGlyT2) was then added. Nonspecific uptake was determined with 10 μM Org24598^27^, (hGlyT1 and mGlyT1 inhibitor), or 5 μM Org25543 (hGlyT2 inhibitor)^54^. The plates were incubated for 15 (hGlyT1) or 30 (mGlyT1 and hGlyT2) minutes with gentle shaking and reactions were stopped by aspiration of the mixture and three times wash with the ice-cold uptake buffer. The cells were lysed, shaken for 3 h and radioactivity was measured by a scintillation counter. The assays were performed in quadruplicate.

### [^3^H]Org24598 binding assay

[^3^H]Org24598 binding experiments were performed in quadruplicate and according to a method previously described^24^. Membranes from CHO cells expressing hGlyT1 and membranes extracted from mouse forebrains (mGlyT1) were used for binding assays. Saturation isotherms were determined by adding [^3^H]Org24598 to mouse forebrain membranes (40 μg/well) and cell membranes (10 μg/well) in a total volume of 500 μL for 3 h at room temperature. Membranes were incubated with 3 nM [^3^H]Org24598 and 10 concentrations of Cmpd1 for 1 h at room temperature. Reactions were terminated by filtering the mixture onto a Unifilter with bonded GF/C filters (PerkinElmer) presoaked in binding buffer containing 50 mM sodium-citrate, pH 6.1 for 1 h and washed three times with the same cold binding buffer. Filtered radioactivity was counted on a scintillation counter. Nonspecific binding was measured in the presence of 10 μM Org24598.

**Extended Data Fig. 1.**
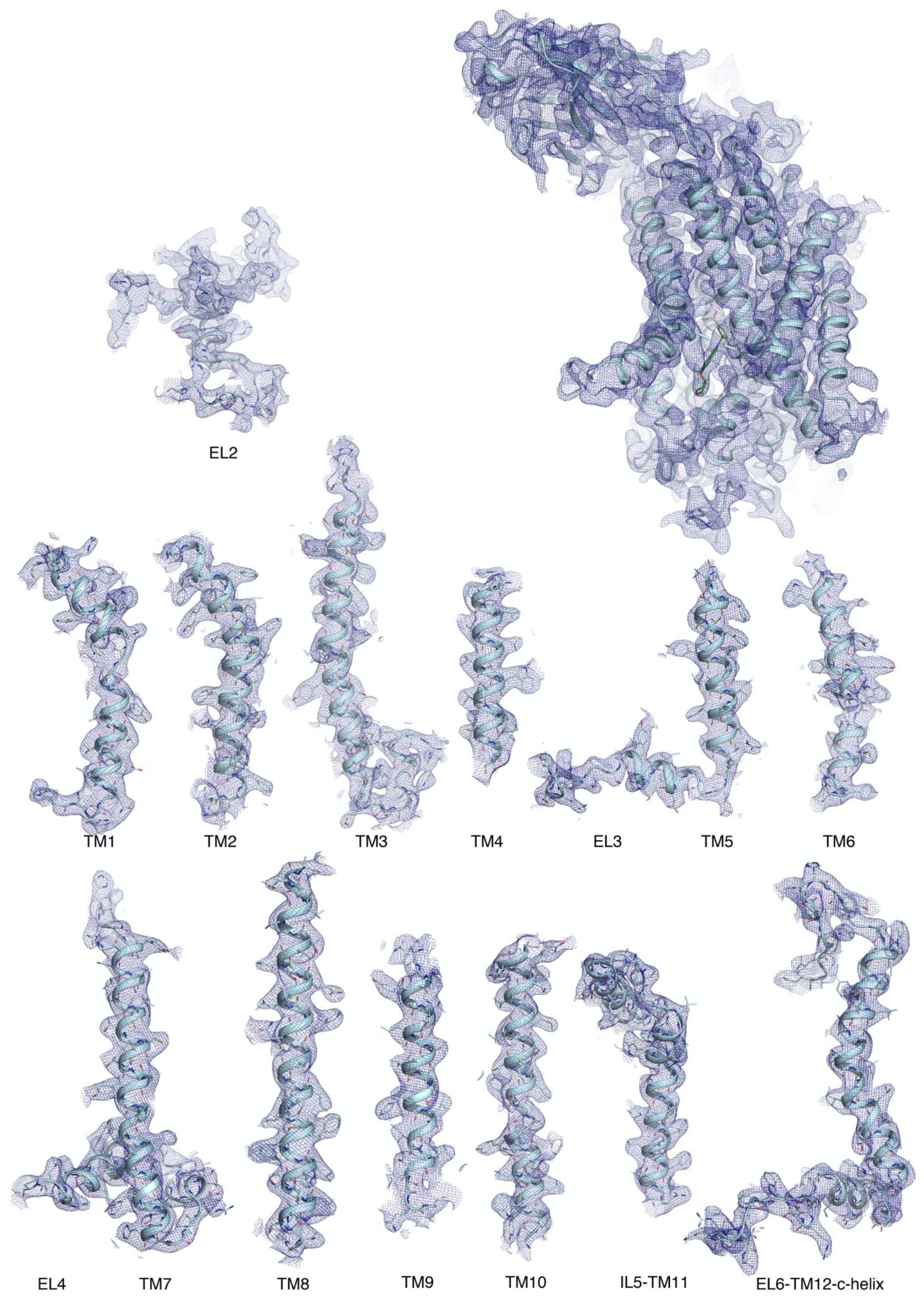
The atomic model of human GlyT1-sybody complex with the bound Cmpd1 inhibitor in the electron density map. The overall structure of GlyT1-sybody complex with bound Cmpd1 (green) and the magnified view of separate transmembrane helices, intra and extracellular loops in 2*F*o – *F*c electron density map (blue) countered at 1.0 r.m.s.d. are shown.

**Extended Data Fig. 2.**
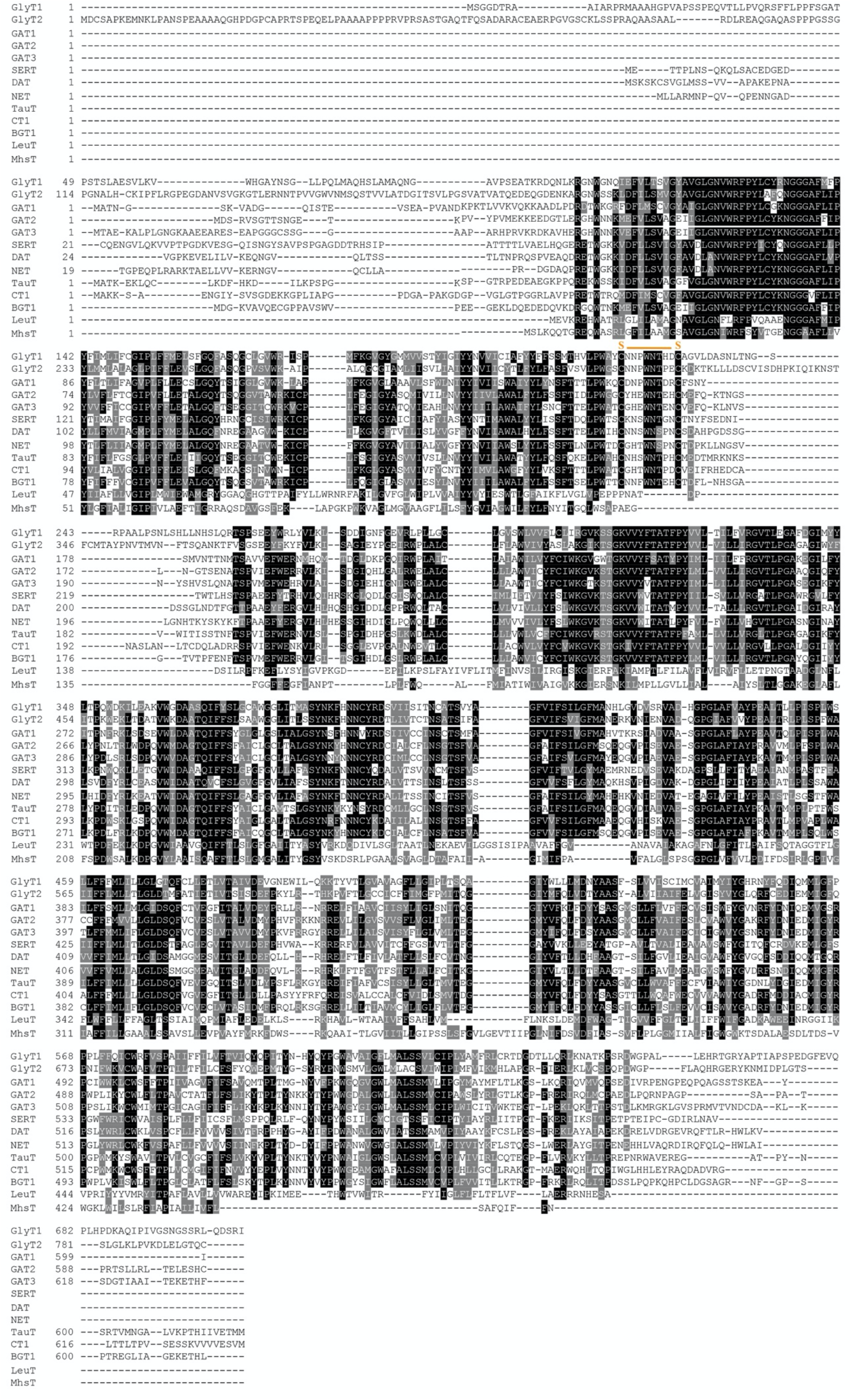
Multiple sequence alignment of NSS family members. Sequence conservation among human NSS family members as well as bacterial homologues, LeuT and MhsT are depicted. Highly conserved residues are highlighted in black.

**Extended Data Fig. 3.**
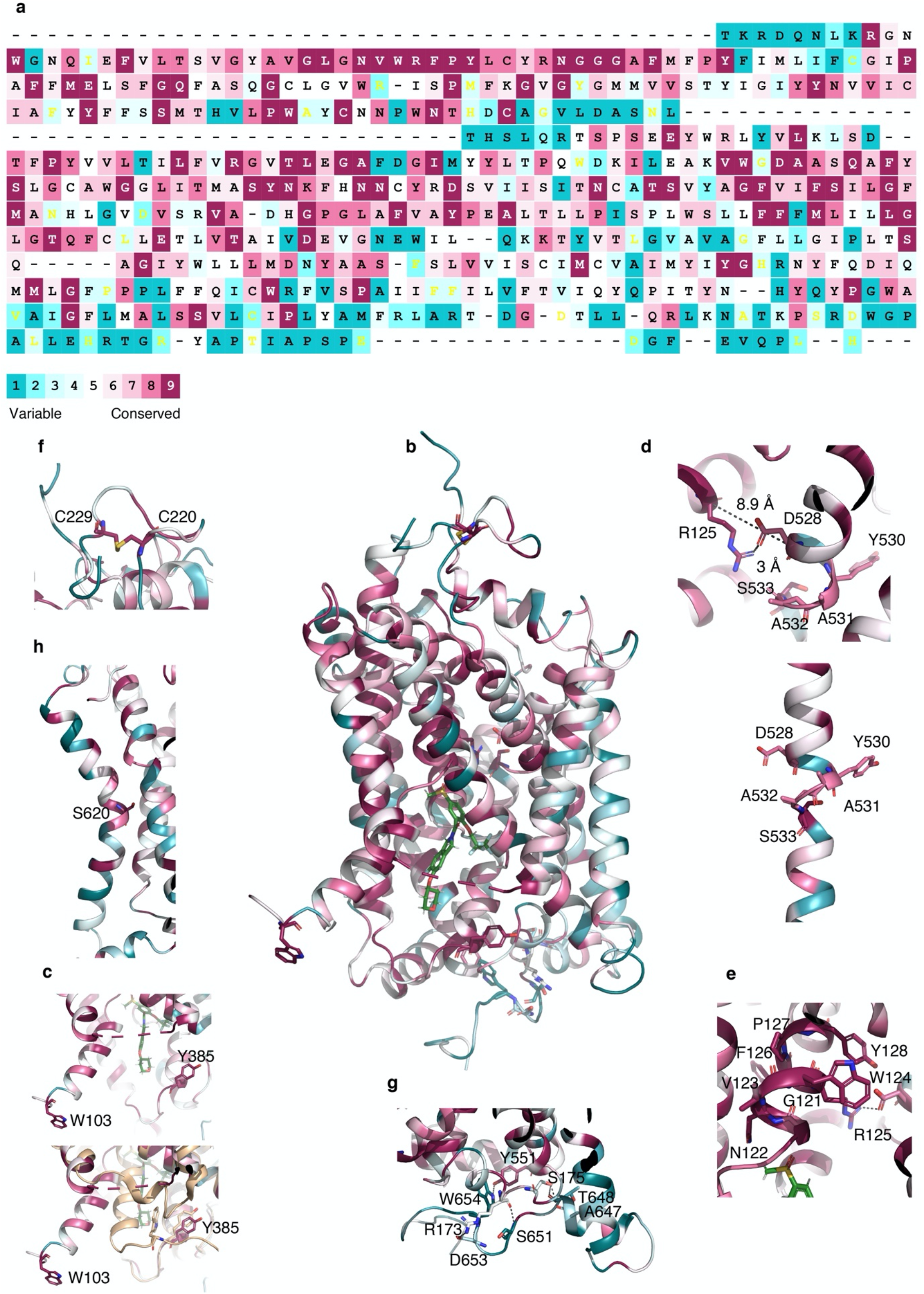
Sequence conservation of hGlyT1. (**a)** The sequence and (**b**) overall structure of human GlyT1 is colored on the basis of ConSurf^55^. (**c**) Disrupted interaction between conserved residues Trp103 (TM1a) and Tyr385 (TM6) due to the hinge-like motion of TM1a in the inward-open structure of hGlyT1 (top) and an overlay of inward facing occluded MhsT (wheat) on inward-open GlyT1 (bottom) is depicted. (**d**) The closed extracellular gate (top) between Asp528 (TM10) and Arg125 (TM1) is depicted. A short non-helical region (bottom) is observed in TM10 at partially conserved Tyr_530_AlaAlaSer_533_ sequence that *supposedly* allows a local flexibility for opening and closing of the extracellular gate between TM10 and TM1. (**e**) The close packing of the extracellular vestibule around Trp124 in the conserved GNVWRFPY motif is shown. (**f**) conserved disulfide bridge on EL2, (**g**) Interacting residues between the C-terminal tail of the transporter and IL1 and IL5 at the intracellular side, and (**h**) Residue Ser620 at the kink of TM12 are shown.

**Extended Data Fig. 4.**
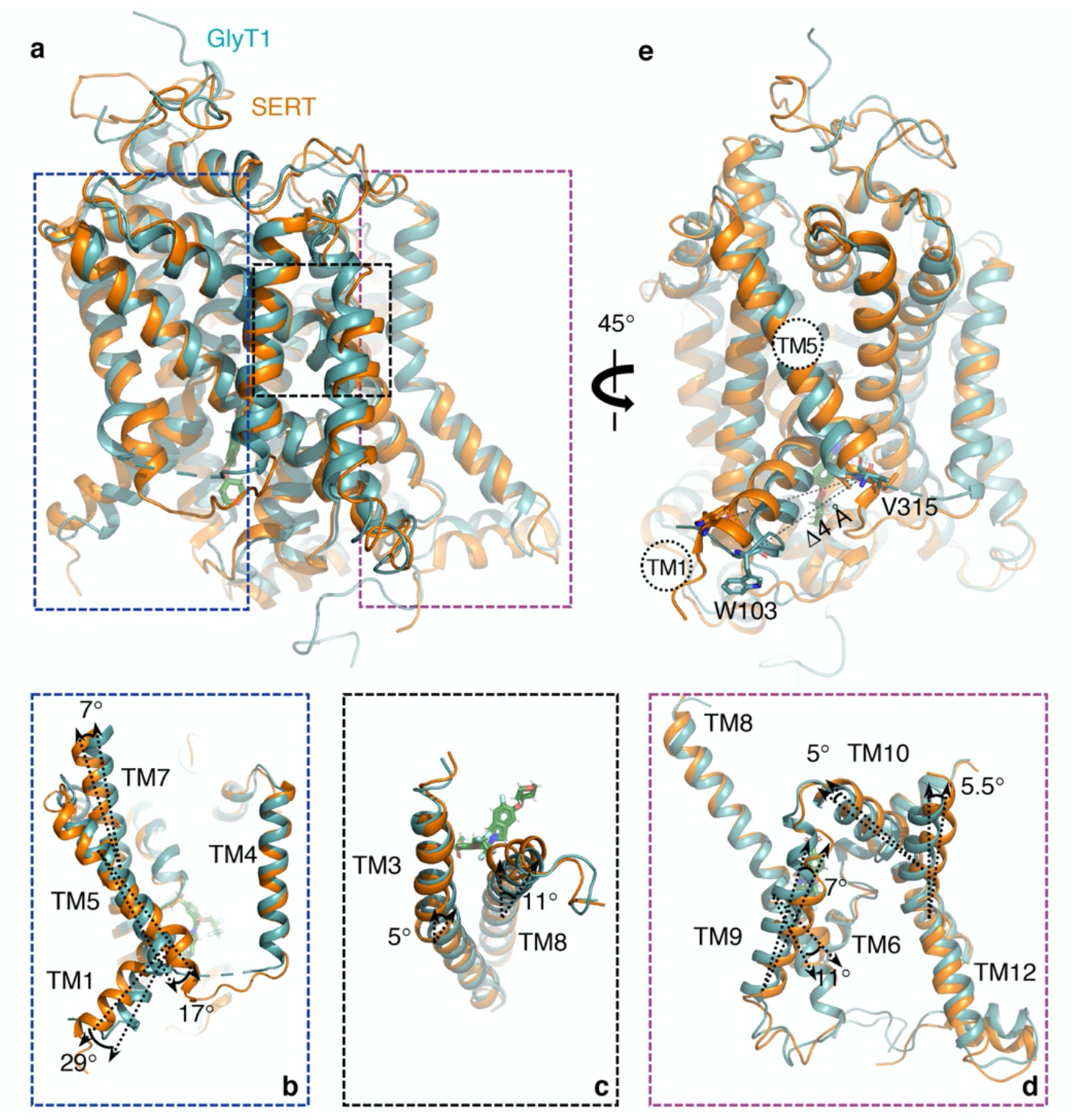
Comparison of inward-open structures of GlyT1 and SERT. (**a**) Secondary structure superposition of GlyT1 (cyan) and SERT (orange) using the so-called scaffold helices TM3-TM4 and TM8-TM9; the TM regions with structural differences are shown in boxes and magnified views are depicted in **b, c**, and **d**. (**b**) The intracellular half of TM1 and extracellular half of TM7 are by 29° and 7°, respectively, closer to the core of GlyT1 compared to the corresponding TMs in inward-open SERT and the intracellular half of TM5 has splayed by 17° further away from the core. (**c**) TM3 in GlyT1 is locally closer to the core halfway across the membrane by 5°. The intracellular half of TM8 have splayed by 11°, further away from the core of GlyT1 compared to SERT. (**d**) On the extracellular side, TM9 is by 7° moved away from TM12, TM10 has shifted by 5° away from TM6 and TM12 is tilted by 5.5° towards the core of GlyT1. The 11° difference at the intracellular half of TM8 is also depicted. (**e**) The intracellular gate to the core of GlyT1 defined by TM1a and TM5 is by 4 Å closer than that of inward-open structure of SERT. C*α* atoms of conserved residues Trp103 of TM1a and Val315 of TM5 were used for the measurement.

**Extended Data Fig. 5.**
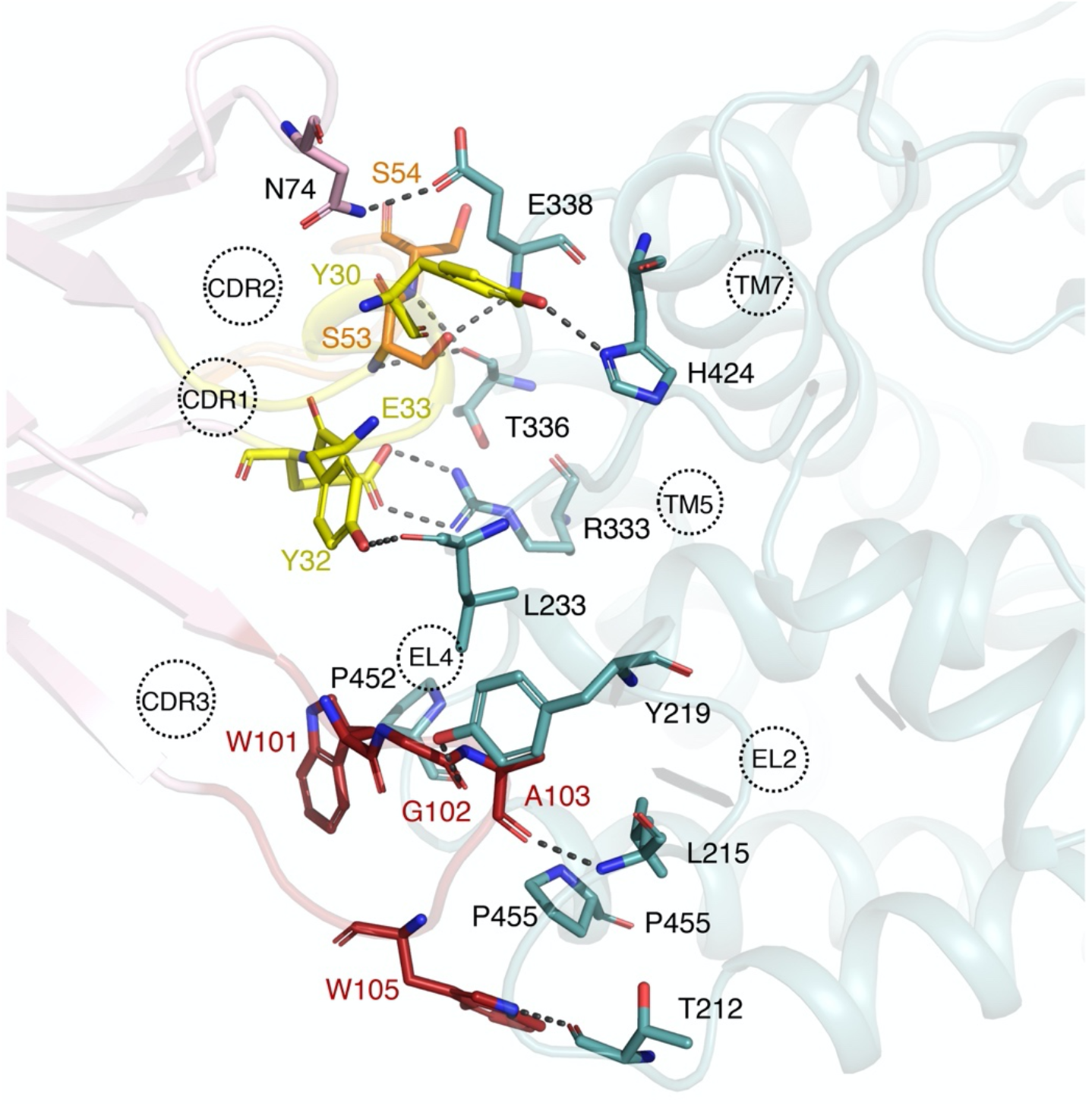
Detailed view of GlyT1-sybody interface. Sb_GlyT1#7 binds to the extracellular segment of GlyT1 through several interactions between the long complementarity-determining region 3 (CDR3), CDR2, and CDR1 of Sb_GlyT1#7 and EL2, EL4, TM5 and TM7 of the transporter. The interface of GlyT1 and sybody was analyzed using contact as a part of CCP4 program suit^56^. Interacting residues of CDR1 (yellow), CDR2 (orange), and CDR3 (red) of the sybody and EL2, EL4 and extracellular ends of TM5 and TM7 of GlyT1 (cyan) are depicted.

**Extended Data Fig. 6.**
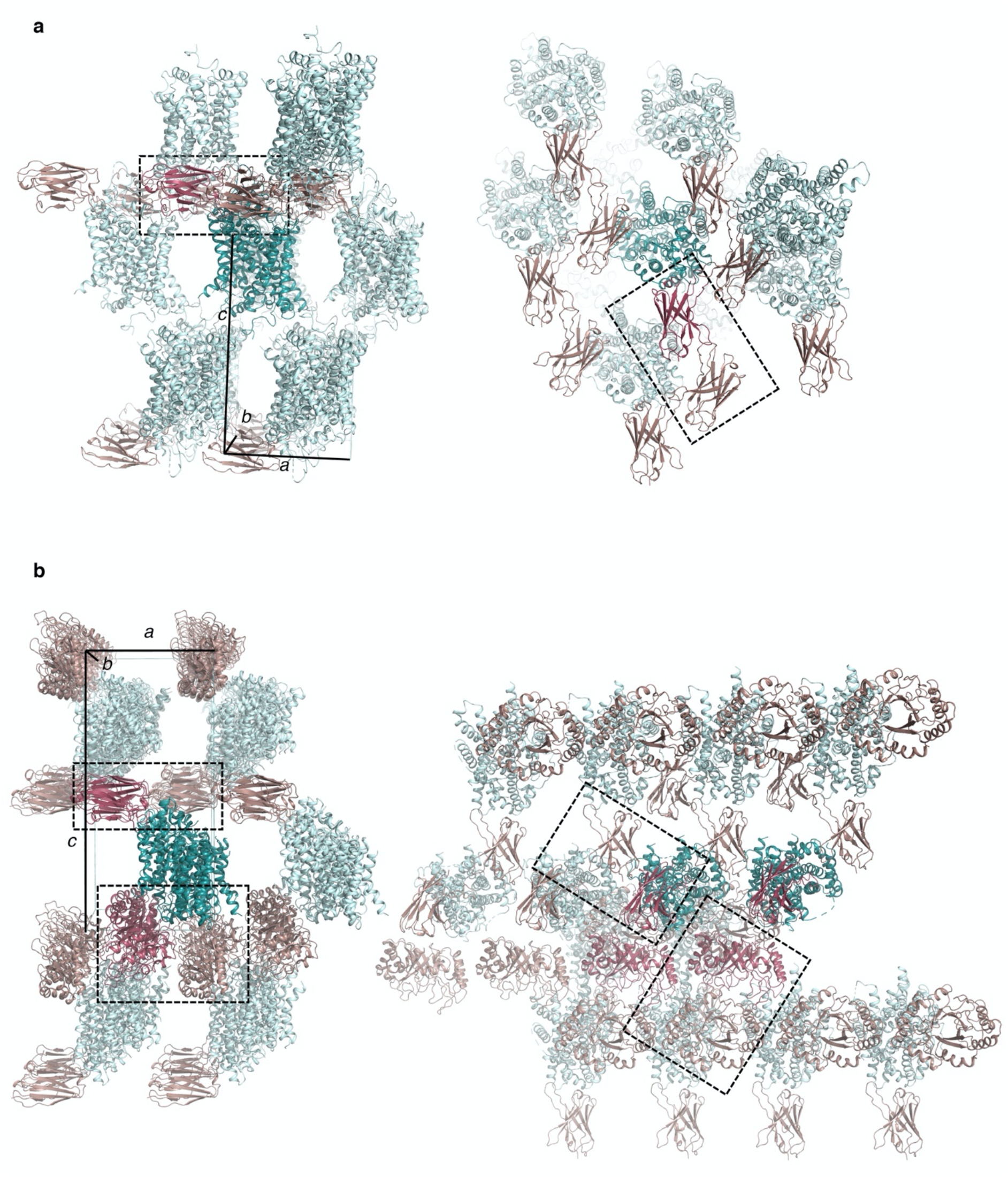
Crystal packing of GlyT1 and GlyT1-Lic. Crystal lattice arrangement viewed from the side and top of GlyT1 (**a**) and GlyT1-Lic (**b**). In GlyT1 crystal contacts exist between adjacent sybodies. In GlyT1-Lic sybodies form the crystal contacts on the extracellular side and adjacent Lichenase fusion proteins on the intracellular side.

**Extended Data Fig. 7.**
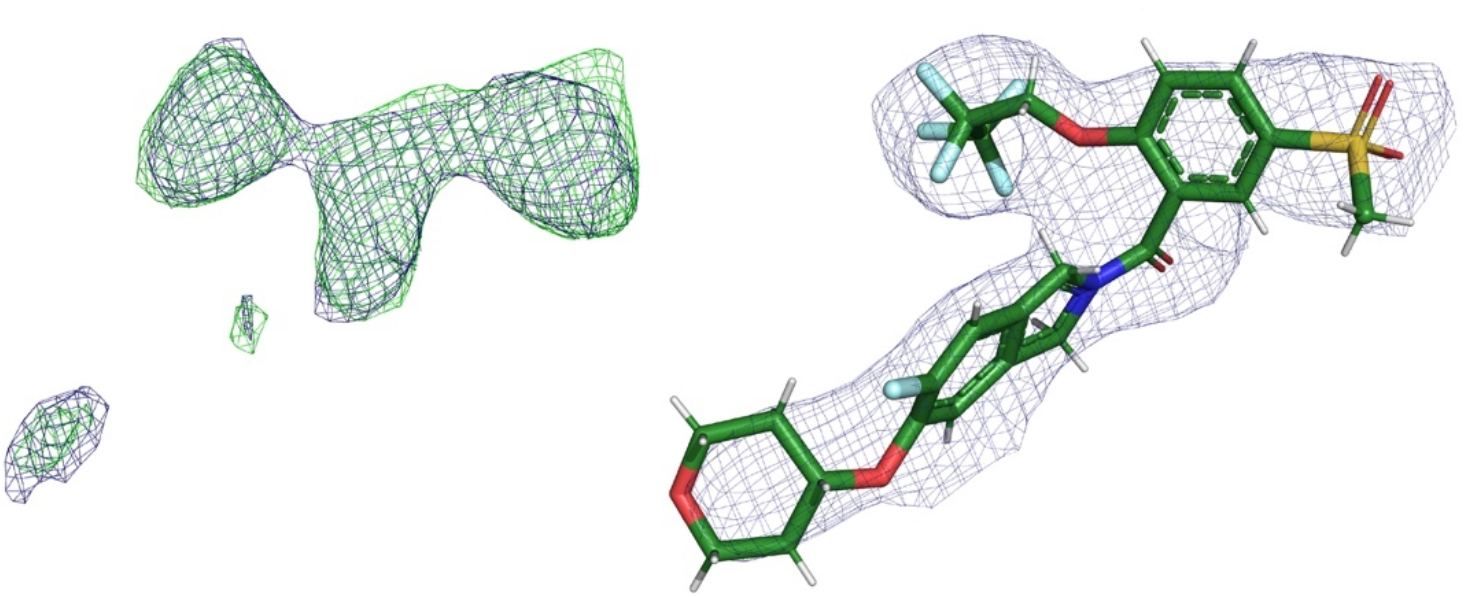
Electron density maps of Cmpd1 inhibitor before and after refinement. The *F*o – *F*c omit (green) and 2*F*o – *F*c (blue) electron density maps of Cmpd1 before placing the inhibitor countered at 3.5 and 1.0 r.m.s.d., respectively is shown on the left and the corresponding 2*F*o – *F*c (blue) electron density map countered at 0.9 r.m.s.d. after placement of the inhibitor and refinement is shown on the right.

**Extended Data Fig. 8.**
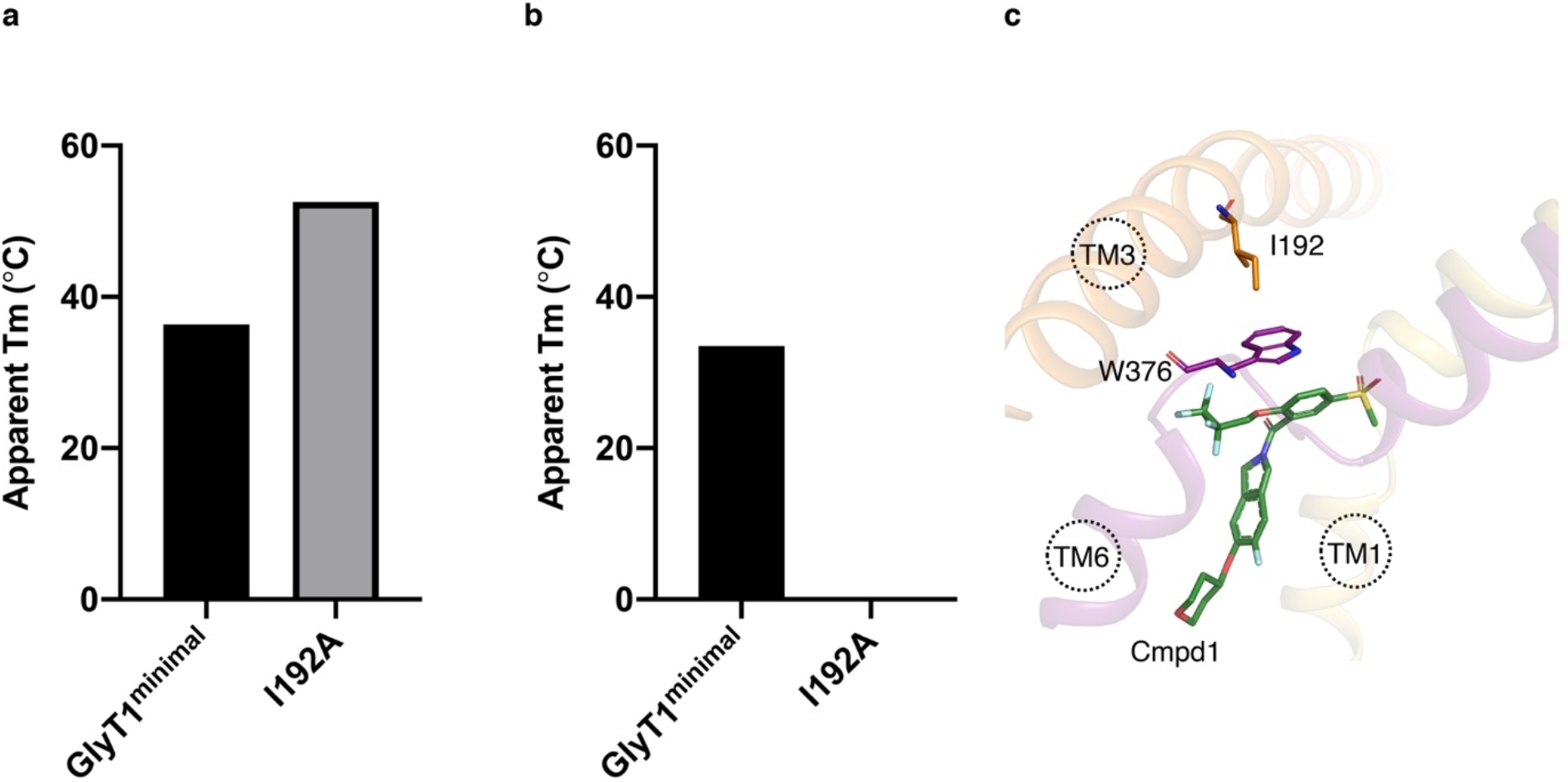
The role of residue Ile192 in inhibitor binding. (**a**) Thermostabilizing effect of Ile192Ala mutation (introduced on GlyT1^minimal^ construct containing also N- and C-termini deletions of residues 1-90 and 685-706, respectively) compared to GlyT1^minimal^ measured by FSEC-TS analysis. (**b**) Non-binding Ile192Ala mutation. SPA-TS analysis shows inability of Ile192Ala mutant to bind the inhibitor Cmpd1. (**c**) Position of Ile192Ala (TM3) stabilizing a rotamer of Trp376 (TM6) in an edge-to-face stacking interaction with inhibitor Cmpd1 is depicted.

**Extended Data Fig. 9.**
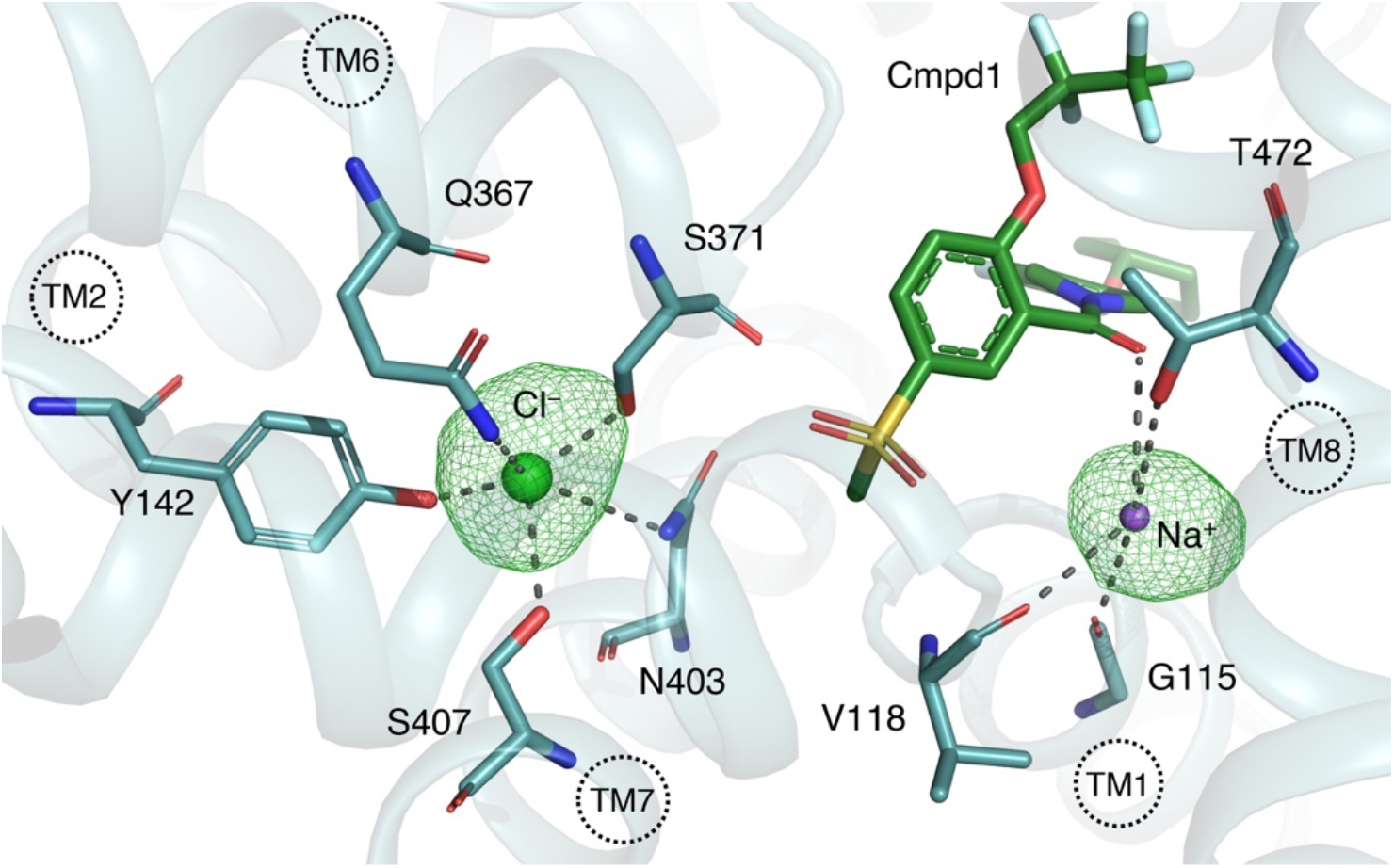
Ion-binding sites. Cl^−^ (light green) and Na^+^ (purple) ions in GlyT1-Lic structure are shown as spheres. The *Fo* − *Fc* simulated annealing^57^ omit maps (green mesh) of Cl^**−**^ and Na^**+**^ ions (a prominent 6.5 r.m.s.d. peak in an unbiased difference map) are shown at 4.0 r.m.s.d. Cmpd1 (green) and residues coordinating Cl^−^ and Na^+^ ions are shown as sticks. The interactions are shown with dashed lines. The chloride ion is coordinated by conserved residues Tyr142 (TM2), Gln367 (TM6), and Ser407 (TM7) similar to the Cl^−^ site in SERT^22^, and further involves Ser371 (unwound region of TM6) and Asn403 (TM7) with a mean coordination distance of 3.0 Å. The sodium ion in Na2 site is within a mean coordination distance of 3.0 Å from the carbonyl oxygen of the conserved residues Gly115, Val118 (TM1) and Thr472 (TM8) as observed in previous structures of NSS transporters as well as the carbonyl oxygen of the Cmpd1 scaffold. The Na1 site as observed in other NSS structures is occupied by the methyl-sulfone substituent of the inhibitor in this structure.

**Extended Data Fig. 10.**
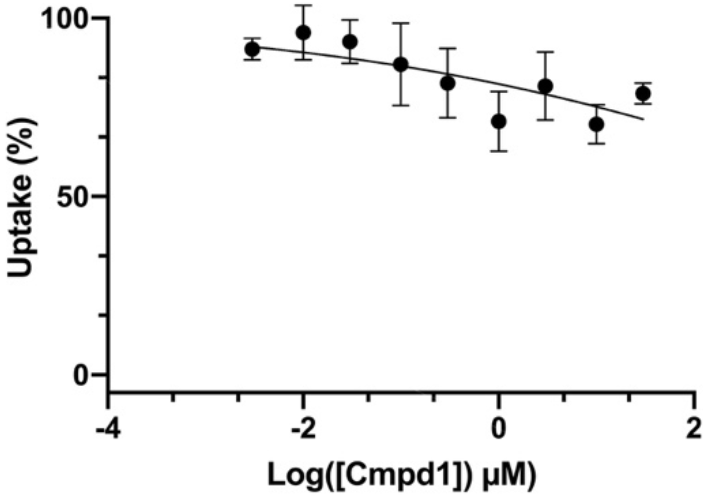
GlyT1 inhibitor, Cmpd1, is not selective against GlyT2. [^3^H]glycine uptake inhibition assay in cells transfected with human GlyT2 cDNA showing that selective inhibitor of GlyT1, Cmpd1, does not inhibit uptake of glycine by GlyT2.

**Extended Data Fig. 11.**
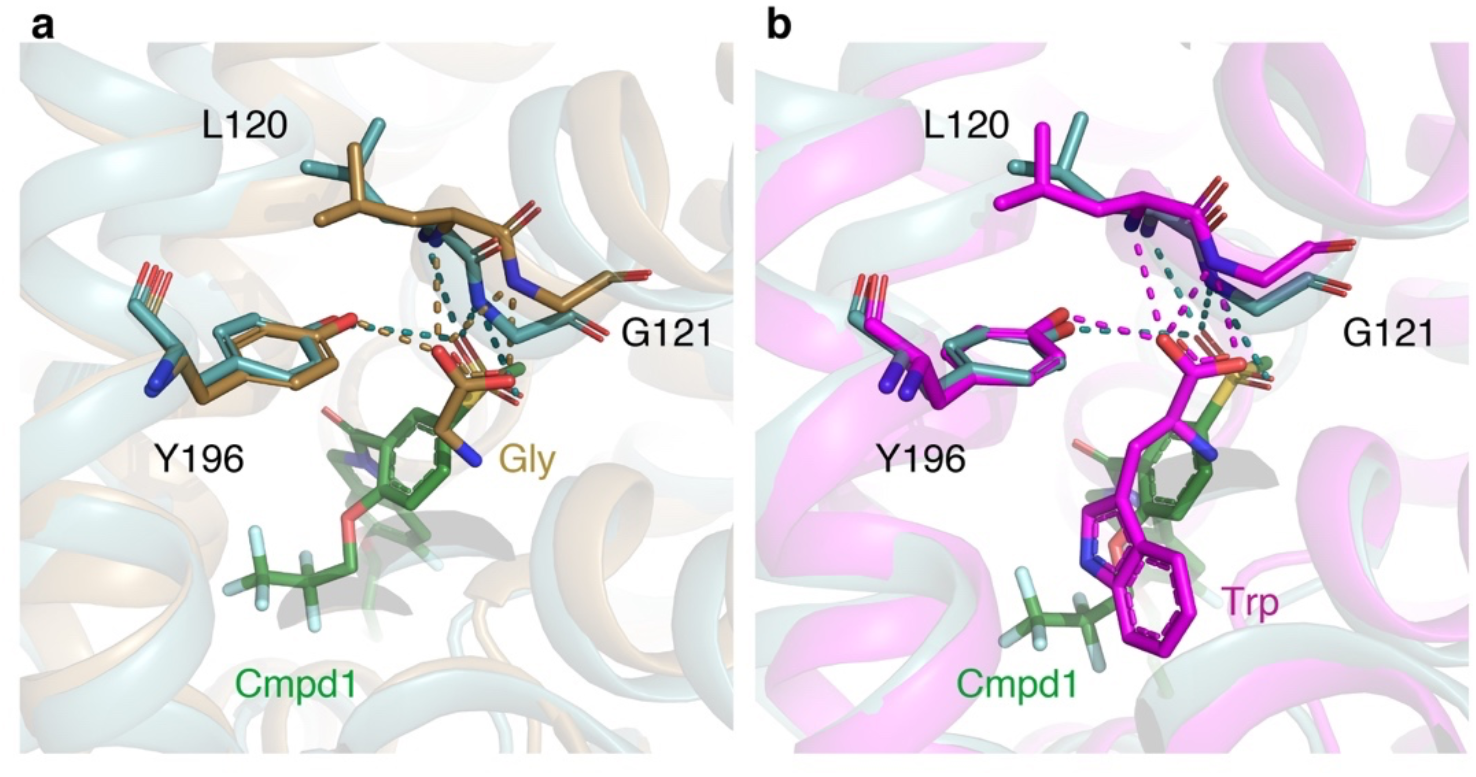
Glycine binding site at GlyT1. (**a**) A superposition of glycine-bound LeuT (sand, PDB ID 3F4J) and (**b**) tryptophan-bound MhsT (magenta, PDB ID 4US3) on Cmpd1-bound GlyT1 (teal) is depicted. The sulfonyl moiety of the inhibitor matches with the carboxylate of glycine or tryptophan. Glycine bound to GlyT1 likely interacts with the backbone amide of Leu120 and Gly121 of TM1 and the hydroxyl group of Tyr196 from TM3 similar to stabilizing interactions in LeuT and MhsT with their respective bound ligands, glycine and tryptophan.

**Extended Data Fig. 12.**
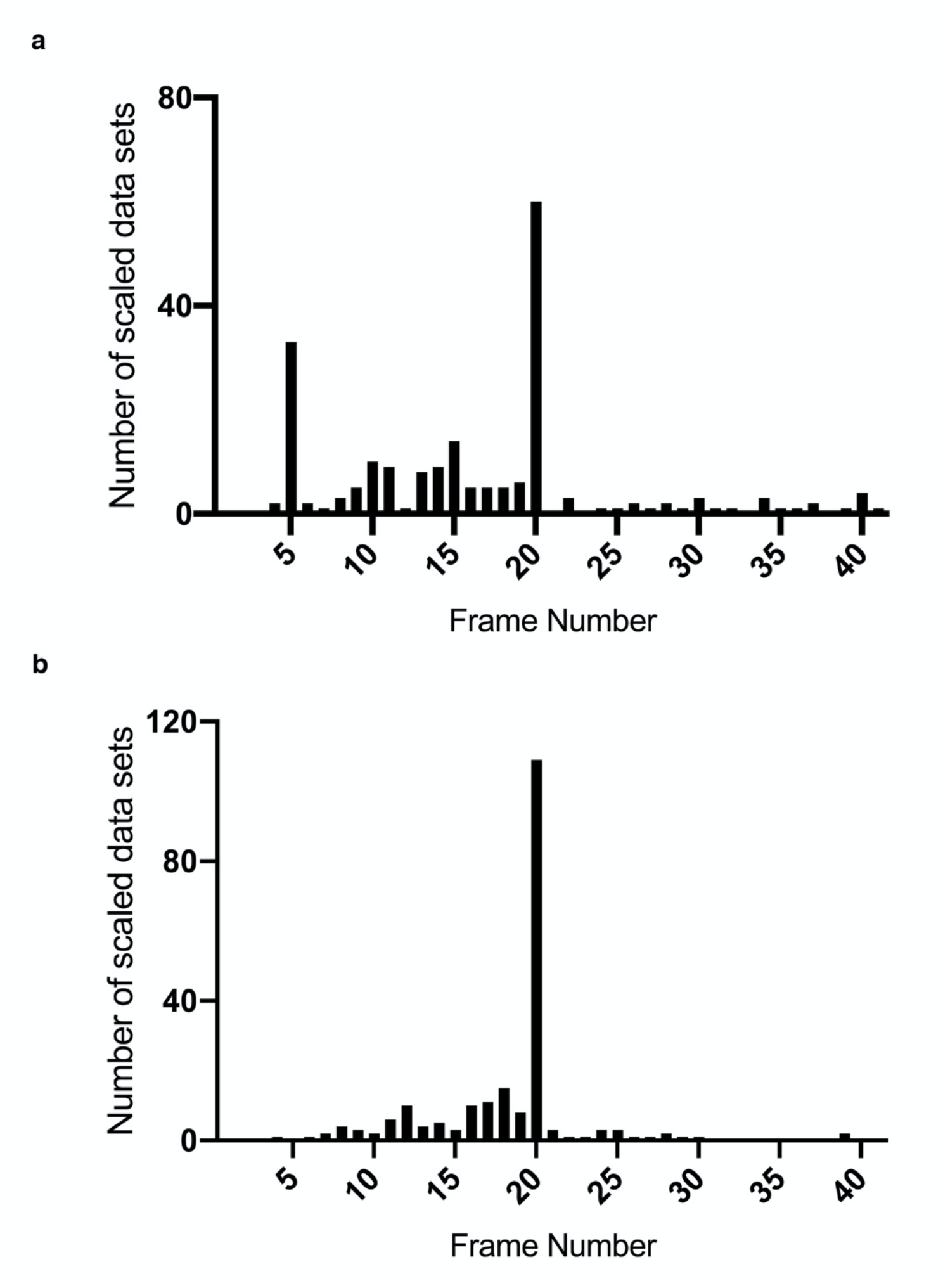
Number of scaled mini data sets per frame for GlyT1 (a) and GlyT1-Lic (b) crystals.

**Extended Data Table 1.** Statistics of unmerged scaled mini data sets of GlyT1 crystals.

| Resolution<br>Limit | Number of Reflections |  |  | Completeness | R-Factor<br>Observed | I/Sigma | R-meas | CC(1/2) |
| --- | --- | --- | --- | --- | --- | --- | --- | --- |
|  | Observed | Unique | Possible |  |  |  |  |  |
| 25.0 | 183 | 30 | 31 | 96.8% | 10.0% | 15.99 | 10.9% | 99.3 |
| 15.0 | 1445 | 221 | 222 | 99.5% | 11.8% | 17.59 | 12.8% | 99.1 |
| 10.0 | 4595 | 670 | 675 | 99.3% | 12.7% | 16.15 | 13.8% | 98.9 |
| 5.0 | 20247 | 3177 | 3211 | 98.9% | 44.6% | 5.19 | 48.6% | 89.3 |
| 4.0 | 46129 | 7219 | 7283 | 99.1% | 61.0% | 4.41 | 66.3% | 84.5 |
| 3.8 | 8249 | 1297 | 1311 | 98.9% | 127.8% | 2.16 | 139.1% | 51.2 |
| 3.7 | 9129 | 1446 | 1445 | 100.0% | 145.5% | 1.74 | 158.3% | 49.3 |
| 3.6 | 9726 | 1582 | 1606 | 98.5% | 153.2% | 1.50 | 167.3% | 46.7 |
| 3.5 | 11305 | 1802 | 1809 | 99.6% | 197.2% | 1.29 | 215.0% | 44.7 |
| 3.4 | 7500 | 1217 | 1227 | 99.2% | 299.2% | 1.00 | 326.4% | 30.1 |
| Total | 153451 | 24071 | 24304 | 99.0% | 41.5% | 4.38 | 45.1% | 97.5 |

**Extended Data Table 2.** Statistics of unmerged scaled mini data sets of GlyT1-Lic crystals.

| Resolution Limit | Number of Reflections |  |  | Completeness | R-Factor Observed | I/Sigma | R-meas | CC(1/2) |
| --- | --- | --- | --- | --- | --- | --- | --- | --- |
|  | Observed | Unique | Possible |  |  |  |  |  |
| 25.0 | 515 | 42 | 42 | 100.0% | 11.9% | 23.98 | 12.5% | 100.0 |
| 15.0 | 4268 | 326 | 327 | 99.7% | 13.9% | 20.11 | 14.5% | 99.8 |
| 10.0 | 13736 | 976 | 975 | 100.0% | 16.9% | 19.27 | 17.6% | 99.5 |
| 8.0 | 18010 | 1291 | 1291 | 100.0% | 23.5 | 13.85 | 24.4% | 99.3 |
| 5.0 | 112156 | 7979 | 7980 | 100.0% | 92.4% | 4.35 | 95.9% | 91.7 |
| 4.8 | 19359 | 1402 | 1402 | 100.0% | 147.0% | 2.79 | 152.7% | 84.2 |
| 4.6 | 22569 | 1618 | 1619 | 99.9% | 177.5% | 2.43 | 184.4% | 79.1 |
| 4.4 | 26270 | 1901 | 1899 | 100.0% | 173.6% | 2.19 | 180.3% | 73.5 |
| 4.2 | 32001 | 2309 | 2309 | 100.0% | 294.4% | 1.48 | 305.7% | 63.4 |
| 4.0 | 37805 | 2784 | 2784 | 100.0% | 460.3% | 1.00 | 478.5% | 42.1 |
| 3.9 | 21006 | 1606 | 1606 | 100.0% | 656.4% | 0.77 | 683.4% | 29.4 |
| Total | 307695 | 22234 | 22296 | 99.7% | 92.1% | 4.42 | 95.7% | 98.5 |

**Extended Data Table 3.** Data collection and refinement statistics.

|  | GlyT1 | GlyT1-Lic |
| --- | --- | --- |
| **Data collection** |  |  |
| Space group | P 1 21 1 | P 1 21 1 |
| Cell dimensions |  |  |
| *a*, *b*, *c* (Å) | 65.17 58.14 122.31 | 116.41 69.71 149.43 |
| $\alpha$, $\beta$, $\gamma$ (°) | 90.00 100.38 90.00 | 90.00 92.86 90.00 |
| Resolution range (Å) | 30 - 3.4 (3.5 – 3.4) | 30 – 3.9 (4.0-3.9) |
| Total reflections | 153292 (14642) | 280831 (26597) |
| Unique reflections | 12641 (1232) | 22219 (2218) |
| R-meas (%) | 44.0 (280.9) | 93.1 (601.4) |
| R-split (%) | 15.1 (69.9) | 28.4 (161.7) |
| R-merge (%) | 42.2 (268.8) | 89.3 (575.5) |
| CC(1/2) | 98.9 (54.1) | 98.4 (26.0) |
| Mean I/σI | 6.57 (1.66) | 4.42 (1.00) |
| Completeness (%) | 99.7 (100.0) | 99.6 (100.0) |
| Multiplicity | 12.1 (11.9) | 13.8 (13.1) |
| **Refinement** |  |  |
| R<sub>work</sub>/R<sub>free</sub> (%) | 21.5/25.8 | 27.7/30.0 |
| No. residues | 659 | 1845 |
| No. atoms | 5258 | 14861 |
| Protein | 5220 | 14781 |
| Ligand/ion | 38/0 | 76/4 |
| Average B-factor | 104.2 | 128.6 |
| Protein | 104.1 | 128.9 |
| Ligands | 120.5 | 76.33 |
| R.m.s. deviations |  |  |
| Bond lengths (Å) | 0.011 | 0.011 |
| Bond angles (°) | 1.67 | 1.55 |
| **Ramachandran plot** |  |  |
| Favored (%) | 95.55 | 94.30 |
| Disallowed (%), (No. residues) | 0.15, (1) | 0.33, (6) |

